# Firing rate diversity lowers the dimension of population covariability in neuronal networks

**DOI:** 10.1101/2024.08.30.610535

**Authors:** Gengshuo John Tian, Ou Zhu, Vinay Shirhatti, Charles M. Greenspon, John E. Downey, David J. Freedman, Brent Doiron

**Affiliations:** Department of Statistics, University of Chicago, Chicago, IL, USA; Grossman Center for Quantitative Biology and Human Behavior, University of Chicago, Chicago, IL, USA; Neuroscience Institute, University of Chicago, Chicago, IL, USA; National Institute for Theory and Mathematics in Biology, Chicago, IL, USA; Center for Living Systems, University of Chicago, Chicago, IL, USA; Department of Neurobiology, University of Chicago, Chicago, IL, USA; Department of Organismal Biology & Anatomy, University of Chicago, Chicago, IL, USA

## Abstract

Neuronal responses are very diverse, with specific stimuli or behaviors eliciting a range of activity across a neuronal population. These responses also show large and correlated trial-to-trial fluctuations that occupy a low-dimensional region in activity space. We link these two aspects of neuronal response in both feedforward and recurrent circuit models and derive the following relation: the more diverse the firing rates of neurons in a population, the lower the effective dimension of their trial-to-trial covariability. We test our prediction using simultaneously recorded neuronal populations from numerous brain areas in mice, non-human primates, and in the motor cortex of human participants. Finally, we show how diverse neural codes lead to better stimulus discrimination, and that when the brain is in a heightened state of processing responses are more diverse and have lower-dimensional fluctuations. In sum, we present an organizing principle of population response that is widely observed across the nervous system and acts to synergistically improve population representation.

## Introduction

Measurement, analysis, and classification of the activity of neuronal populations and how it represents sensory, cognitive, and motor variables is essential to uncover the computational goals of the nervous system [1]. The full space of neuronal responses is extremely complex, so researchers need to focus on certain metrics of population response to attain theoretical understanding of any neural code.

One prominent feature of population response is the broad distribution of firing rates among neurons when responding to a stimulus or behavior [2–4]. For a given condition, some neurons fire at high rates while most of the remaining neurons fire at low rates, creating a diverse response profile. This aspect of neural representation has been extensively studied in the framework of sparse coding, demonstrating the wide range of computational advantages it brings such as disentanglement and energy efficiency [5].

Another important feature of neuronal response is variability, where population activity can vary drastically across identical trials of a task [6–8]. The structure of neuronal variability underlies behavior [9], and is thus essential for understanding the computations leading to task successes and failures. Critically, trial-to-trial fluctuations are shared across the neuronal population and are often concentrated around a low-dimensional subspace [10–16]. It has been shown that restricting noise to directions orthogonal to the informative subspace facilitates linear decoding downstream [17, 18].

While trial-averaged responses and any fluctuations about the average are formally statistically unrelated quantities, they are connected in neuronal networks through the particular cellular and circuit mechanisms that generate neuronal activity patterns [19]. In our study, we consider both the diversity of firing rates and the dimensionality of population covariability in circuit models. Using techniques from random matrix theory, we derive our central result: increased diversity of the mean response across neurons lowers the effective dimension of population covariability. Our prediction is verified with electrophysiology datasets from multiple brain areas in mice, non-human primates, and human participants. We use our theory to show that increased firing rate diversity and the associated low-dimensional population variability act synergistically to improve population decoding. This prediction is supported by experimental evidence where neurons exhibit both more diversity in their firing rates and a lower-dimensional variability in brain states with higher information processing demands.

### The effective dimension of population covariability decreases with firing rate diversity

Consider a population of *N* neurons with output response vector *r* ∈ ℝ^*N*^ which is a random variable with mean *R* (represented as a diagonal matrix for notational consistency) and covariance matrix *C* (Fig. 1a). We quantify the uniformity of the mean response *R* by the Treves-Rolls measure *D*(*R*) (Fig 1a,b) [20–22]. A *D*(*R*) near 0 marks a diverse mean response distribution where a few neurons fire at much higher rates than the rest and a *D*(*R*) near 1 suggests neurons fire at rates similar to one another. We measure the effective dimension of population covariability with the participation ratio *D*(*C*) (Fig 1a,b) [23, 24], where a *D*(*C*) near 0 indicates the concentration of fluctuations around a small number of dimensions and a *D*(*C*) near 1 implies that fluctuations are distributed more or less uniformly across different directions and are thus high-dimensional. While the functional forms of *D*(*R*) and *D*(*C*) are identical, they nevertheless measure different aspects of neuronal response, namely the location of the mean versus the shape of the variability (Fig. 1b; Methods).

**Fig. 1:**
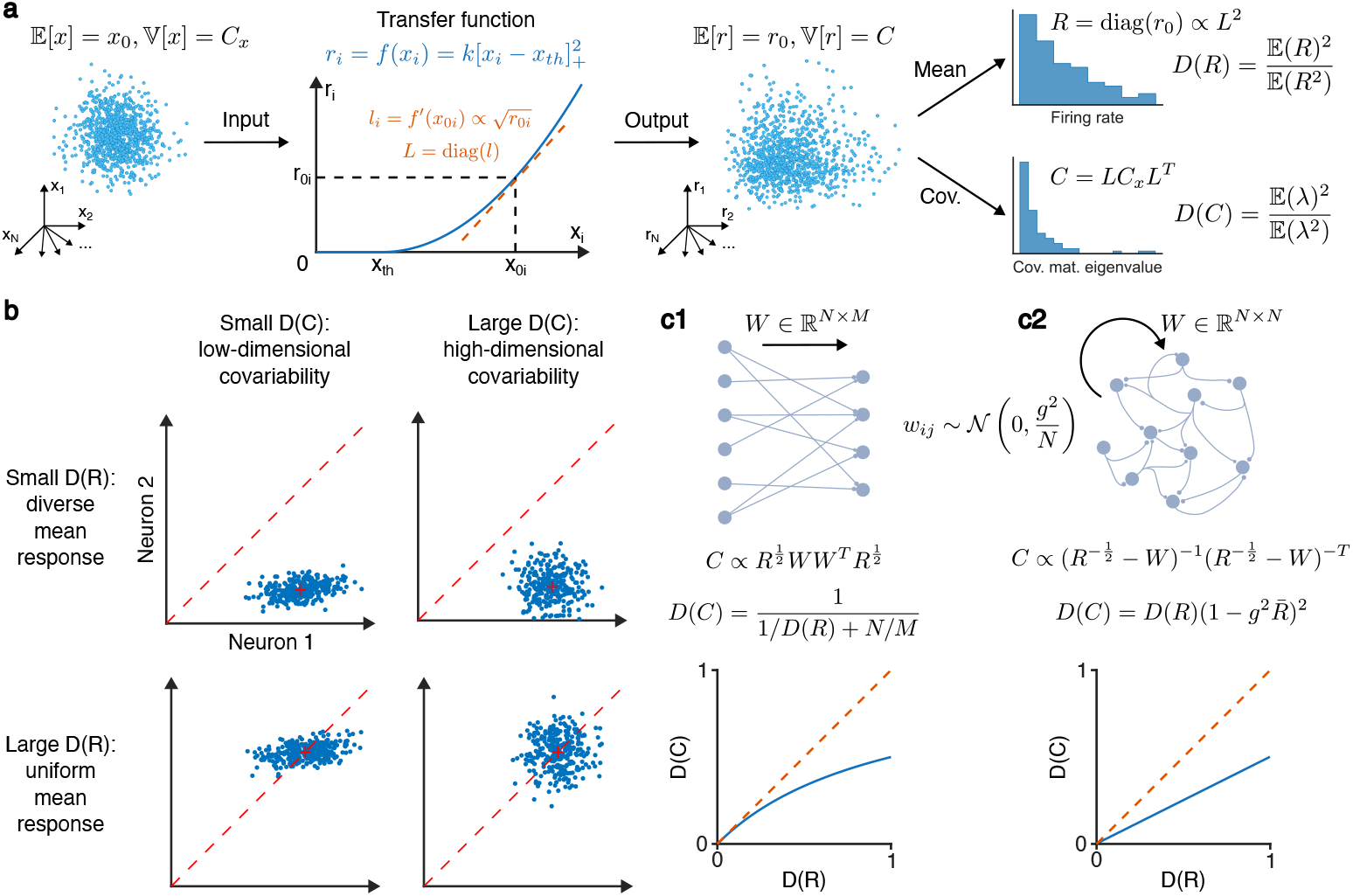
Relating firing rate diversity and the effective dimension of population covariability. (**a**) The distribution of the synaptic input *x* and that of the spiking output *r* are related through the nonlinear transfer function of neurons *f*, which determines the correspondence between the firing rate *r*_*i*_ of a neuron and its gain *l*_*i*_. This leads to a relation between the mean and covariance of *r*, and in turn one between the firing rate uniformity *D*(*R*) and the effective dimension *D*(*C*). (**b**) Illustration of the distribution of neuronal response in the spike count space with small/large *D*(*R*)/*D*(*C*), respectively. Each dot is the response of a neuron pair for a given realization of input, and the location and shape of the cloud of responses determine *D*(*R*) and *D*(*C*), respectively. (**c**) Two ways of shaping the covariance matrix. Left: Feedforward projection *W* from a population of size *M* in the previous layer to the current layer of size *N* leads to a positive but sublinear relation between *D*(*C*) and *D*(*R*) (Eq. (2)). Right: Recurrent connectivity *W* in a population of size *N* leads to a positive linear relation between *D*(*C*) and *D*(*R*) (Eq. (4)).

While *R* and *C* are unrelated to each other from a statistical standpoint, there is a mechanistic argument that connects them. Consider the vector of inputs received by the neurons *x* ∈ ℝ^*N*^. For each neuron *i*, its output *r*_*i*_ is generated by passing the input *x*_*i*_ it receives through the transfer function *f*, i.e., *r*_*i*_ = *f* (*x*_*i*_). Let the mean and covariance matrix of *x* be *x*_0_ and *C*_*x*_. Linearization yields the relation *C* = *LC*_*x*_*L*^*T*^ between input and output covariances, where *L* is the diagonal matrix of neuronal gains *l*_*i*_ = *f* ^*′*^(*x*_0*i*_) (Fig. 1a). Intuitively, the gain describes how sensitive a neuron’s output is to changes in its input, and the diagonal form of *L* describes the neuron-wise modulation of population variability by the gain. When the transfer function *f* is nonlinear, the gain *l*_*i*_ depends on the linearization point *x*_0*i*_ and thus the firing rate *r*_0*i*_ = *f* (*x*_0*i*_). Experiments have shown that the input-output curve of neurons can be well described by a threshold-power law function [25]:

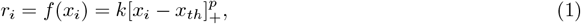

where [·]_+_ = max(·, 0), *x*_*th*_ is the firing threshold, *k* is a constant scaling factor, and *p* is the power of the transfer function. Following previous work, we set the power *p* to be 2 unless otherwise stated [26], which would imply that the neuronal gain is associated with the firing rate by *L*^2^ ∝ *R*.

In our framework the firing rate distribution *R* affects the covariance matrix *C* through the neuronal gain *L*, but the exact relationship between *D*(*C*) and *D*(*R*) depends on the specific form of the input covariance *C*_*x*_. In the simplest case where *C*_*x*_ = *I*, we have *C* = *R* and *D*(*C*) = *D*(*R*), meaning that a more diverse mean response produces covariability with a lower effective dimension. It is unclear whether this trend holds for input covariances with more complex structures. To explore this question, we take a mechanistic approach and consider how *C*_*x*_ is generated in feedforward and recurrently coupled circuits.

#### Feedforward circuit

First, let us consider a neuronal population that receives input from another population of size *M* through feedforward connections *W* (Fig. 1c1). Assuming that the activity of the upstream layer is i.i.d., the structure of *C*_*x*_ arises completely from *W* and *C*_*x*_ = *WW*^*T*^ (this is without loss of generality, as any upstream covariance structure can be absorbed into *W* and we arrive at the same expression). We model *W* by drawing its elements independently from a Gaussian distribution, which is a typical setup in circuit models [24, 27]. The problem then reduces to deriving the eigenspectrum of *C* from those of *C*_*x*_ and *R*, both of which are known. Critically, classical scalar approaches in probability theory fail when we consider two random matrices (*C*_*x*_ and *R*) because of the non-commutative nature of matrices (i.e., *C*_*x*_*R* ≠ *RC*_*x*_). This issue can be addressed by adopting techniques from free probability as applied to random matrix theory [28]. This allows us to derive the full eigenvalue distribution of *C* (Supplementary Information B.2; Fig. S1a). From this we calculate the moments of *C*’s eigenspectrum, which in turn gives us its effective dimension (Supplementary Information B.3; Fig. 1c1):

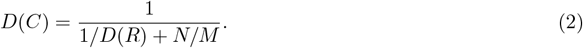

We see that the positive relation between *D*(*C*) and *D*(*R*) when *C*_*x*_ = *I* persists in the feedforward model (Fig. 1c1, bottom). However, for moderate *N/M* we have *D*(*C*) *< D*(*R*), implying that coupling acts to lower the effective dimension. The intuition is that a realization of the random wiring will promote joint activity along certain directions, while suppressing it along other directions. Any deviation from isotropic joint fluctuations acts to lower the dimension of *C*.

#### Recurrent circuit

A key assumption in the derivations for the feedforward model is the free independence between the input covariance *C*_*x*_ and the firing rates *R*. A sufficient condition for free independence is the independence between *C*_*x*_ and *R* combined with rotational invariance of the distribution of *C*_*x*_ [28], both of which are satisfied in the feedforward setting. In a recurrent network, however, the input to each neuron is heavily influenced by the activity of other neurons in the population, leading to a dependence of *C*_*x*_ on *R*. We thus need to reexamine the eigenspectrum of the output covariance *C* in a recurrent circuit.

Let *W* now denote the recurrent connectivity in the network, with elements drawn independently from a Gaussian distribution with mean 0 and variance *g*^2^*/N* (Fig. 1c2) [29, 30]. It is a standard result [29–34] that linearizing the network activity about an operating point gives us the following expression for the covariance matrix (Supplementary Information C.1):

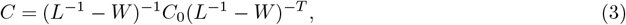

where *C*_0_ is the covariance matrix of the *external* input noise. We assume for now that the external input noise has the same magnitude across neurons and are uncorrelated (*C*_0_ = *I*). In this case the expression for *C* can no longer be cleanly separated into the input covariance and gain in the form of *C* = *LC*_*x*_*L*^*T*^, demonstrating the fundamentally recurrent nature of the network interactions.

If neurons are uncoupled (*g* = 0) and have uniform gain (*L* = *I*), we have the simple case of *C* = *I* and *D*(*C*) = 1. This is the largest achievable effective dimension, where joint activity fluctuates equally in all directions and is independent of the mean firing rates of the neurons. If we keep the neurons uncoupled but allow for diverse gain from nonlinear transfer functions (Eq. (1)) and a wide distribution of firing rates as before, we again have *C* = *R* and *D*(*C*) = *D*(*R*) with firing rate diversity lowering the effective dimension of covariability. Previous works have studied the case where connectivity is heterogeneous (*g >* 0) but the neuronal gain remain homogeneous (*L* = *I*). It was derived that *D*(*C*) = (1 − *g*^2^)^2^, showing that the effective dimension is lower for more diverse coupling as quantified by a larger *g* [29, 30, 33].

The above two scenarios each incorporate one aspect of diversity in the network (connectivity and gain), with each having a critical impact on the effective dimension of response variability. The natural question to ask is then what happens when both of these factors are present in the system and interacting in a nontrivial manner, as is the case for real neuronal circuits. We again turn to (operator-valued) free probability theory to derive the eigenvalue distribution of the covariance matrix *C* from those of *W* and *R* [28] (Supplementary Information C.2; Fig. S1b). This allowed us to calculate the moments of *C*’s eigenvalue distribution, which then yields our central analytic result (Supplementary Information C.3; Fig. 1c2):

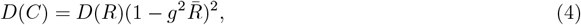

where 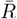 is the population mean firing rate. We see that diversity in the mean response (smaller *D*(*R*)) and the connectivity (larger *g*) combine multiplicatively to lower the effective dimension of population covariability (smaller *D*(*C*)). Intuitively, the heterogeneities in neuronal gains and in synaptic wiring are preserved in the recurrent dynamics and both contribute to the anisotropicity of the population fluctuations. Critically, *D*(*C*) only depends on the first two moments of the firing rates *R* irrespective of its exact distribution. This was verified in simulations using Eq. (3) (Fig. S2a). We remark that Dahmen et al. [30] approached the problem by directly modeling a linear recurrent network and treating *W* as the *effective* connectivity matrix which includes both synaptic strengths and neuronal gains. Their approach interprets the *g* parameter not as measuring the variance of connection weights but as the spectral radius of *W* . However, their method cannot properly treat the heterogeneity in gains, at least not in its current form (see Supplementary Information C.6 for a detailed discussion).

Finally, we can combine the feedforward and recurrent models by allowing the external input covariance matrix *C*_0_ in Eq. (3) to not be the identity matrix while still requiring it to be freely independent from *W* and *R*. We then obtain a general formula for *D*(*C*) (Eq. (S12)), where the relation between *D*(*C*) and *D*(*R*) is still positive but no longer linear (but instead sublinear, similar to Fig. 1c1), and both a smaller *D*(*C*_0_) and a larger *g* lead to smaller *D*(*C*). That is, heterogeneity in recurrent connectivity, low dimensionality of the external input, and diversity of firing rates all contribute to a smaller effective dimension of population covariability. The feedforward model is recovered as a special case by setting the recurrent weights to 0 and equating *C*_*x*_ with *C*_0_.

In total, in randomly coupled networks of neurons with nonlinear transfer (i.e., *L* depends on *R*) there is a positive relation between the uniformity of firing rates *D*(*R*) and the effective dimension of covariability *D*(*C*). Before we test this result in a variety of cortical datasets, we explore how circuit structure can affect the [*D*(*R*), *D*(*C*)] relation.

#### Breaking the relation between *D*(*R*) and *D*(*C*) in structured networks

Recall that the sufficient condition we used to guarantee free independence requires both independence and rotational invariance of one of the matrices involved [28]. In the feedforward model, the input covariance *C*_*x*_ (or *C*_0_ in the more general setting of Eq. (3) and Eq. (S12)) is rotationally invariant; in the recurrent model, the recurrent weight *W* is rotationally invariant. Intuitively, rotational invariance can be understood as the joint distribution of eigenvectors having no directional bias in the vector space nor any relation with the eigenvalues. This assumption can be broken when there is significant structure in the distribution of *W* or *C*_0_ beyond i.i.d. elements.

Indeed, real neuronal circuits almost never fit the description of an unstructured graph. For example, there is often spatial organization in the circuit’s wiring, where neurons that are close to one another (in either physical or feature selectivity space) are more likely to connect to each other and receive correlated inputs (due to, e.g., common feedforward and feedback projections) [35, 36]. We explored the effect of structured wiring on the [*D*(*R*), *D*(*C*)] relation by simulating a network where neurons are randomly placed in a two-dimensional space and where *W* and *C*_0_ are generated according to the distance between neurons (Fig. 2a,b; Methods). The correlation between *D*(*C*) and *D*(*R*) decreases as input correlations become more widespread (Fig. 2d) or recurrent connections become more spatially concentrated (Fig. 2e), both of which moves the system further away from the free independence assumption. The measured correlation may also vary depending on which neurons are sampled and what conditions are explored in the simulated recording session, forming a distribution of correlation values (Fig. 2f). We therefore see that the [*D*(*R*), *D*(*C*)] relation is not trivially guaranteed to hold in neural data, as in many circuits we expect structure in both input correlations and recurrent wiring.

**Fig. 2:**
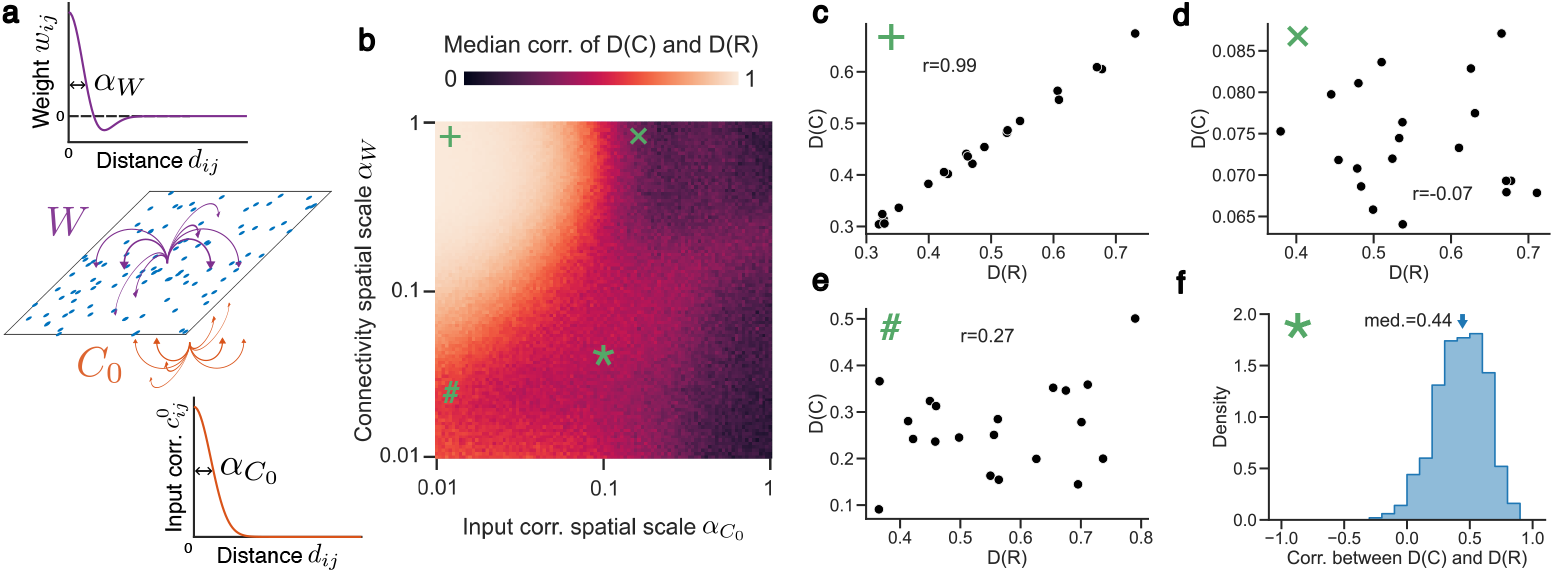
Spatially dependent connectivity *W* and input noise covariance *C*_0_ can break the [*D*(*R*).*D*(*C*)] relation. (**a**) Neurons are randomly placed on a plane with those close to each other having both larger connection weights and external input correlations. The parameters *α*_*W*_ and 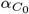 quantify the widths of the connectivity profile and the input correlation profile respectively. The simulated population size is 100. (**b**) Heatmap of the [*D*(*R*), *D*(*C*)] correlation as *α*_*W*_ and 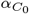 change. Note that the axes are in log scale. (**c**) Example [*D*(*R*), *D*(*C*)] scatter plot in a regime with high [*D*(*R*), *D*(*C*)] correlation. Note that a larger *α*_*W*_ and a smaller 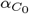 signify *W* and *C*_0_ being closer to the rotational invariance assumption in our theory. (**d**) Example [*D*(*R*), *D*(*C*)] scatter plot in a regime with low [*D*(*R*), *D*(*C*)] correlation due to structure in recurrent connectivity. (**e**) Example [*D*(*R*), *D*(*C*)] scatter plot in a regime with low [*D*(*R*), *D*(*C*)] correlation due to structure in external input. (**f**) Example distribution of the [*D*(*R*), *D*(*C*)] correlation across realizations of *W* and *C*_0_ with moderate spatial structure in the network.

### Validation in neural data across species and brain areas

We set out to test the positive relationship between the effective dimension of population covariability *D*(*C*) and the uniformity of firing rates *D*(*R*) in neural data. We remark that it is generally difficult to disentangle the effects due to external input versus recurrent interactions, as only the recorded neurons’ output activity is accessible to the experimenter, and other information such as connection weights or direct measurements of neuronal gain are hard to obtain. In our theory, both feedforward and recurrent effects may modulate the quantitative formula for *D*(*C*), and it is likely that both are present in real neuronal circuits, but since their qualitative predictions on the positive relationship between *D*(*C*) and *D*(*R*) converge, we remain agnostic to which specific model underlies the observations in data. However, for convenience, we shall use Eq. (4) as the reference model in the following since it presents a linear relationship that is easier to interpret.

To test how *D*(*C*) varies with *D*(*R*), one needs to sample a range of *D*(*R*) values. This can be achieved by presenting different experimental conditions such as distinct sensory stimuli, task contexts, and cognitive states, all of which modify the firing rate distribution of the recorded population. In a single experimental session, it is usually reasonable to assume that the macroscopic statistics of synaptic connectivity do not change appreciably, and a stable relation between *D*(*C*) and *D*(*R*) should be observed. We demonstrate this pipeline with simulations, where we generated covariance matrices using Eq. (3) with the firing rates *R* drawn from a log-normal distribution ℒ𝒩 (*µ, σ*). We varied *D*(*R*) by changing *µ* and *σ* while keeping *g* and 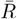 fixed and plotted the resulting [*D*(*R*), *D*(*C*)] pairs (Fig. 3a; Methods). Indeed, the simulation result follows theoretical predictions closely.

**Fig. 3:**
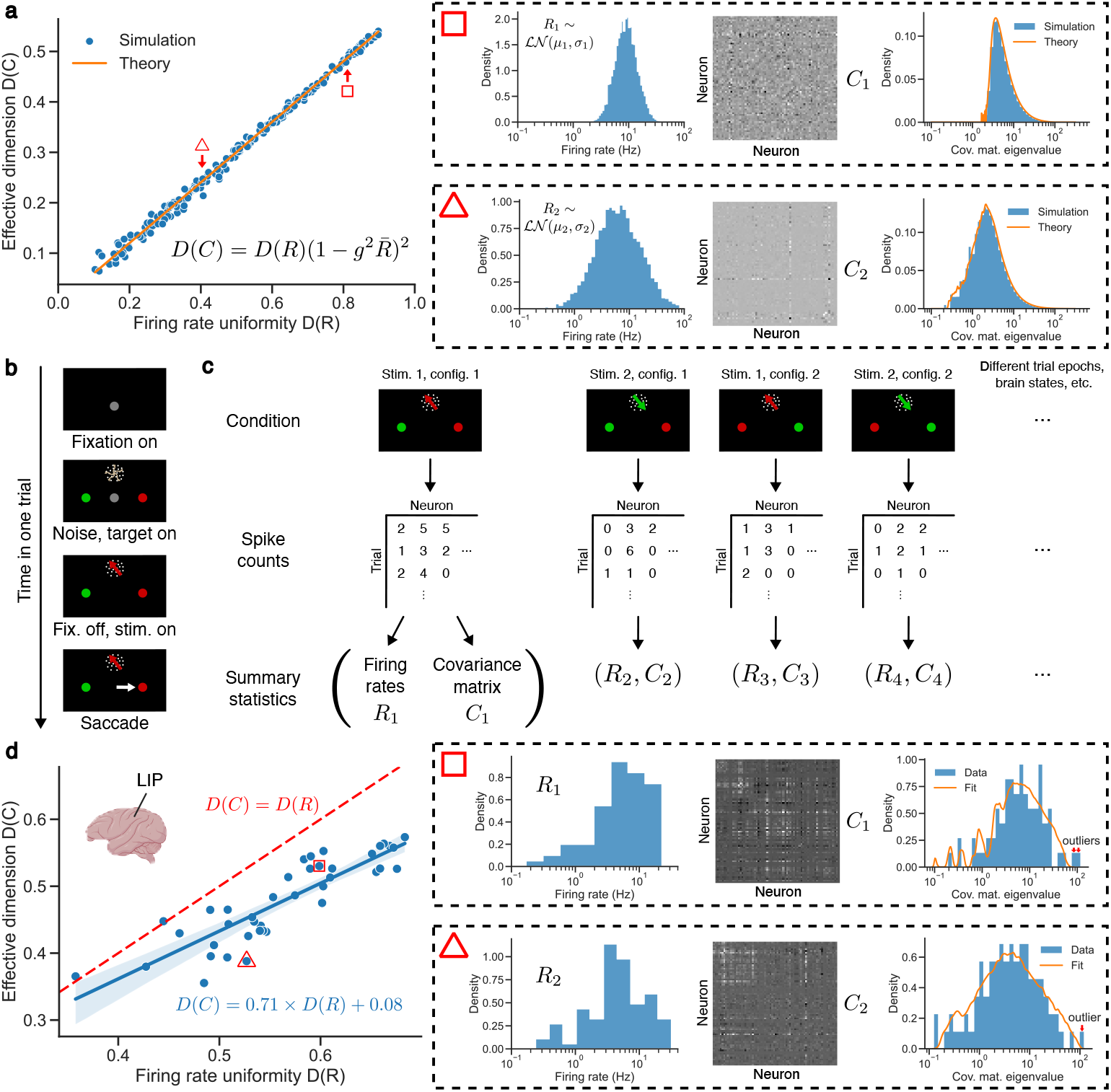
Validation of our theory in population recordings in area LIP in monkeys. (**a**) Theoretical prediction of the effective dimension *D*(*C*) as firing rate uniformity *D*(*R*) changes (Eq. (4)) matches simulation results. The population size is *N* = 5, 000. *g* = 0.15 and 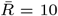. The boxes on the right demonstrate how a point in the plot is produced. Each point in the left plot has a different firing rate distribution *R* drawn from a log-normal distribution with parameters designed to keep the mean firing rate the same but covers a range of *D*(*R*) values. The sampled *R* vector is used to generate the covariance matrix *C* according to Eq. (3) while keeping the connectivity *W* unchanged. It can be directly observed that the triangle point (bottom right) with smaller *D*(*R*) and *D*(*C*) has wider firing rate distribution and covariance matrix eigenvalue distribution than that from the square point (top right). The rightmost column additionally shows that the eigenvalue distribution predicted by our theory (Eq. (S9)) matches the simulation. (**b**) Structure of the visual categorization task (see Methods for details). (**c**) Different experimental conditions set the system at different operating points with different firing rates *R* and covariance matrices *C*, which are calculated from the spike count data matrix produced by each condition. Different conditions here could mean different experimental variables such as stimulus parameters (here the direction of moving dots) and response target configurations (here whether to saccade to the left or right target for the red category). They could also mean different epochs in the task as shown in (b) or different brain states such as spontaneous versus evoked. (**d**) The linear relationship between *D*(*C*) and *D*(*R*) verified with monkey LIP data. Linear regression: slope=0.71, intercept=0.08, Pearson correlation coefficient (PCC) *r* = 0.87, *p* = 1.8 × 10^−13^. The number of recorded neurons is *N* = 88. The red dashed line is the identity line *D*(*C*) = *D*(*R*). Similar to (a), the boxes on the right illustrate how each point in the plot on the left corresponds to a different (*R, C*) pair as conditions vary. The rightmost column demonstrates how fitting our model to the data identifies outlier eigenvalues.

To explore our prediction in real cortical population response, we trained monkeys to perform a visual categorization task (modified from previous experiments [37], see Methods; Fig. 3b) and recorded spike train data from the lateral intraparietal cortex (LIP). Different experimental conditions set the recorded neuronal populations at different operating points, producing many pairs of mean responses *R* and covariance matrices *C* (Fig. 3c; Methods), which mirrors what took place in the simulations (Fig. 3a). As discussed earlier, it can be assumed that the overall connectivity statistics remain largely unchanged within the time frame of one experimental session. Further, the population mean firing rate 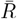 does not widely fluctuate across conditions (Fig. S4a; also see Fig. S4b for a demonstration in a larger dataset). We can then plot the effective dimension *D*(*C*) against firing rate uniformity *D*(*R*) across conditions (Fig. 3d) and see that they are indeed positively related, as predicted by our theory (Fig. 1c).

We note that the mathematical technique we used to derive the eigenvalue distribution of the covariance matrix *C* can only address the “bulk” of the distribution (meaning the portion that has positive measure as the system size grows to infinity). Finite-rank perturbations to either the connectivity matrix *W* or the input noise covariance matrix *C*_0_ would result in a finite number of outliers in *C*’s eigenvalues, which is beyond the scope of our theory [29] (Supplementary Information C.5). Since these perturbations originate from factors that cannot be measured experimentally, we need to exclude them in order to test our theory. We achieve this by fitting our model to the data to obtain a reference model of *C*’s eigenvalue distribution. We can then statistically test which eigenvalues cannot be explained by this reference model, and identify them as outliers that likely originate from low-rank perturbations to the system [29, 30] (Fig. 3d; Methods). Interestingly, as we shall see later, in many datasets the relationship between *D*(*C*) and *D*(*R*) persists even without removing the outliers (Fig. S6).

One might be tempted to use the slope of the linear fit to the [*D*(*R*), *D*(*C*)] scatter plot to infer the variance parameter *g* of the recorded circuit’s synaptic connections. We would caution against this because the slope of the fit is affected by multiple factors other than *g*. First, as we have seen in the feedforward model, the value of input covariance dimensionality *D*(*C*_0_) may significantly modulate the [*D*(*R*), *D*(*C*)] relationship. Second, the recorded neurons constitute only a small sample of the underlying population, and it can be shown (Supplementary Information D.2) that under this spatial sub-sampling, *D*(*C*) and *D*(*R*) are still linearly related in the recurrent model (Eq. (S20), Fig. S2d), but the coefficient of this proportionality depends on the activity and connectivity of the unobserved population and can vary considerably across sessions that record from different subpopulations. Previous work has also shown that the proportion of excitatory and inhibitory neurons sampled in the recording has significant impact on the population activity structure including its dimensionality [38].

A third confounding factor is that the actual input-output transfer functions in the recorded population is unknown, and may even be heterogeneous across neurons. To probe how it affects our theory, we reexamine different values of the power *p* in Eq. (1) (Eq. (4) was developed under *p* = 2). We have analytic formulae for general *p*’s (Eq. (S11)), where *D*(*C*) and *D*(*R*) are again positively related, but the relation is no longer linear (Fig. S3a). Simulations show that *D*(*C*) and *D*(*R*) are positively correlated as long as the gain depends on the firing rate (*p* ≠ 1; but see Supplementary Information C.4) (Fig. S3b). We also explored the effect of having a distribution of *p*’s across neurons and found that this additional source of diversity lowers *D*(*C*) (Fig. S3c), but again preserves the positive correlation between *D*(*C*) and *D*(*R*) (Fig. S3d). Taken together, we see that while the [*D*(*R*), *D*(*C*)] relationship is largely intact, the slope of a linear fit would clearly depend on the distribution of *p* in the population, again preventing us from estimating the connectivity parameter *g* from data.

Since our theory was developed for a generic recurrent circuit, it could in principle be applied more generally across diverse brain areas and species. We analyzed data from various cortical and subcortical brain areas across monkeys (Fig. 4a-c), mice (Fig. 4f-h), and human participants (Fig. 4k, Fig. S5) including both novel data we collected (monkey areas LIP, FEF, and SC, and human M1; see Methods) and previously published datasets [39–42, 44]. The positive correlation between *D*(*C*) and *D*(*R*) overwhelmingly holds across all the data we examined when compared to shuffled data (Fig. 4d, i; Methods). At the same time, the wide distribution of correlation values may reflect the moderate levels of structure present in local circuits (compare with Fig. 2f).

**Fig. 4:**
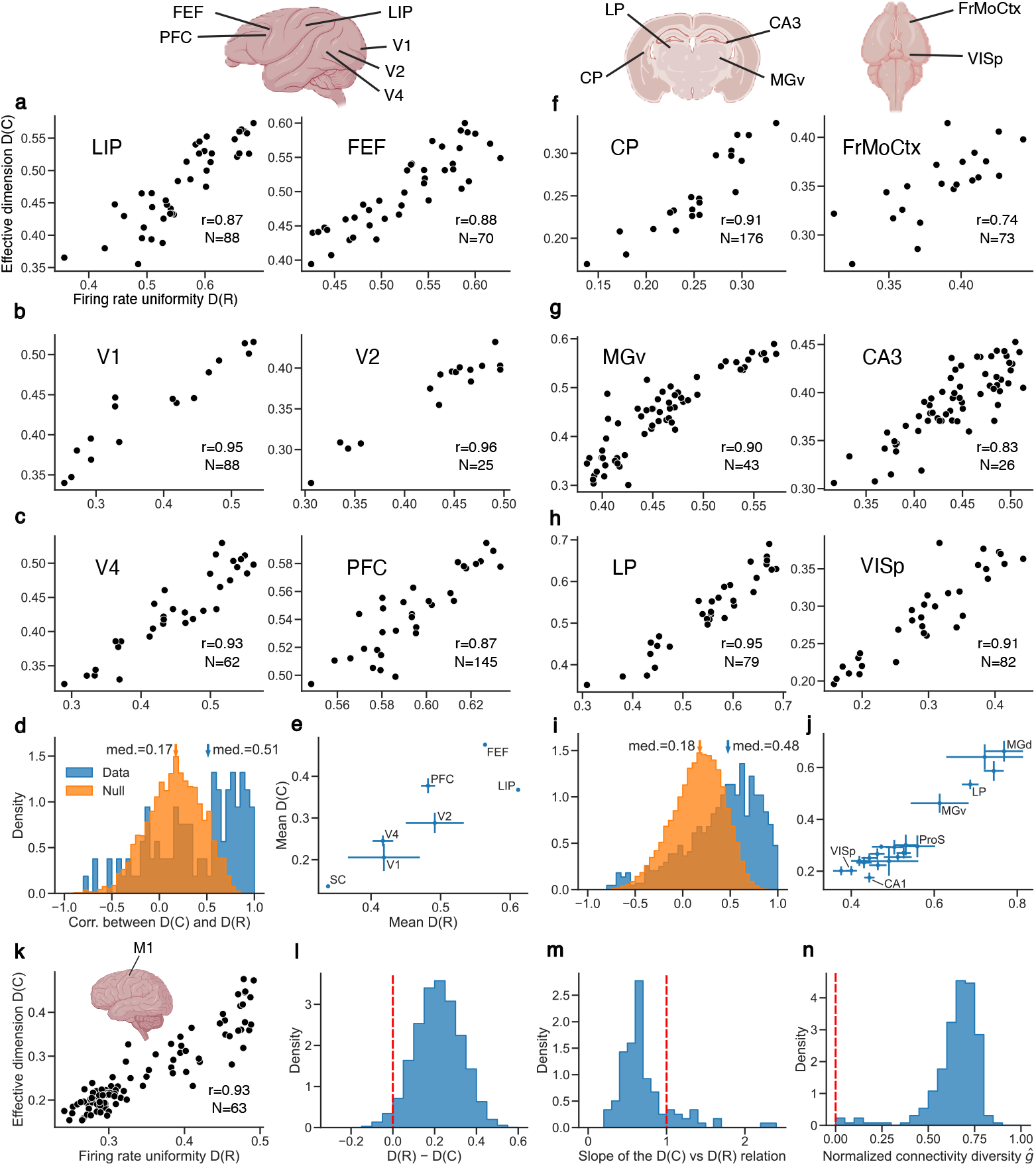
The relationship between *D*(*C*) and *D*(*R*) verified in electrophysiology datasets from multiple brain areas in monkey (a-e), mice (f-j), and human participants (k). Each scatter plot (a-c, f-h, k) shows data from one recorded population in a single session across a range of conditions as in Fig. 3d. *r* is the Pearson correlation coefficient between *D*(*C*) and *D*(*R*), and *N* is the number of recorded neurons. (**a**) Data recorded from monkey LIP (lateral intraparietal) and FEF (frontal eye field) cortices (see Methods). (**b**) Monkey visual areas V1 and V2 data taken from [39]. (**c**) Monkey visual area V4 and prefrontal cortex (PFC) data taken from [40]. (**d**) Aggregate *r* values from all monkey datasets in (a-c) compared to shuffled data. 7 brain areas and 32 sessions in total. The data distribution is significantly larger than the null (median test, *p* = 1.1 × 10^−4^). (**e**) Mean *D*(*C*) and *D*(*R*) across conditions for each brain area in the monkey datasets. Error bars are standard errors of the mean across sessions. To show the original geometry of the neuronal response, outlier eigenvalues were not removed here. (**f**) Mouse caudoputamen (CP) and frontal motor cortex (FrMoCtx) data taken from [41]. (**g**) Mouse medial geniculate nucleus, ventral part (MGv) and CA3 data taken from [42, 43]. (**h**) Mouse lateral posterior nucleus of the thalamus (LP) and primary visual area (VISp) data taken from [44]. (**i**) Same as (d) but for mouse datasets in (f-h). 40 brain areas and 79 sessions in total. The data distribution is significantly larger than the null (median test, *p* = 7.2 × 10^−50^). (**j**) Same as (e), but for the mouse dataset [42]. (**k**) Data recorded from human M1. (**l-n**) Recurrent connectivity has a large impact on *D*(*C*). The histograms are over all brain areas and all sessions in dataset [42]. The red dashed lines are what would be expected for independent Poisson neurons, which are far away from the data distributions in blue. (**l**) Distribution of *D*(*R*) − *D*(*C*). (**m**) Distribution of the slope obtained by performing linear regression on *D*(*C*) against *D*(*R*). Only data where the correlation between *D*(*C*) and *D*(*R*) are greater than 0.5 is included because the slope is not very informative otherwise. (**n**) The distribution of the normalized connectivity diversity parameter *ĝ* obtained from fitting our model to covariance matrix eigenvalue distributions in data.

As mentioned earlier, even when we do not remove the outlier eigenvalues when calculating *D*(*C*), the correlation between *D*(*C*) and *D*(*R*) often (but not always) remains high (Fig. S6), depending on whether these outliers are large enough and are correlated with *D*(*R*) in a way that negates the change in the bulk. Another notable observation is that, when we plot the *D*(*C*) and *D*(*R*) averaged across conditions for each brain area, they are still strongly correlated (Fig. 4e, j), even though each area presumably has very different connectivity structures.

One might argue that the positive correlation between *D*(*R*) and *D*(*C*) is expected if we simply consider the network activity to be a perturbation from a population of uncoupled and uncorrelated Poisson-like neurons with a uniform Fano factor, where *D*(*C*) = *D*(*R*). To refute this argument we provide four lines of evidence. First, the distribution of pairwise noise correlations suggests neurons are far from uncorrelated (Fig. S7a). Second, if a network was a small perturbation from a set of uncoupled neurons, then its output covariance matrix *C* would be a perturbation of a diagonal matrix, and each eigenvector would be dominated by one neuron. This is not the case in our neural data (Fig. S7b). Third, uncoupled Poisson-like neurons would produce *D*(*C*) = *D*(*R*), but we almost always observe *D*(*C*) values that are smaller than their corresponding *D*(*R*) values (Fig. 4l). In line with this, the variations of *D*(*C*) across conditions are also smaller than those of *D*(*R*) as indicated by the slope of the [*D*(*R*), *D*(*C*)] plot being smaller than 1 (Fig. 4m). Finally, our theory in the recurrent model requires that 0 ≤ *ĝ <* 1, where 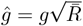 is the normalized connectivity diversity parameter (Supplementary Information C.2). A perturbation from an uncoupled network would have *ĝ* near zero. However, the *ĝ* fitted from data is typically larger than 0.5 (Fig. 4n), suggesting that recurrent connections (or feedforward connections and external input covariance) play an important role in generating neuronal variability. Indeed, our model always explains the eigenvalue distribution in the data much better than an uncoupled model (Fig. S7c-d).

In sum, these results show that the positive [*D*(*R*), *D*(*C*)] relationship largely holds for neuronal pop-ulations across the brain, suggesting that the structures in their recurrent wiring *W* and input covariance *C*_0_ are typically not prominent enough to break our theoretical prediction. However, these analyses were all restricted to data from single brain regions. It is unclear whether this relationship still holds when neurons from multiple brain regions are put together, where there is presumably significant block structure in *W* . We examine this in the next section.

### Combining multiple brain regions breaks the relationship between *D*(*R*) and *D*(*C*)

Multi-regional brain connectomes show a clear block structure, with distinct within-area and between-area connectivity [45]. Simulations show that the correlation between *D*(*C*) and *D*(*R*) can be degraded at the multi-population level by such block structures, but remain largely intact for the activity in each population separately (Fig. S8). To investigate how such interareal connectivity structures affect the [*D*(*R*), *D*(*C*)] relation, we analyzed data recorded simultaneously from monkey V4 and PFC [40] (Fig. 5a1) as well as from mouse VISp and VISl [42] (Fig. 5b1). When we pool neurons from two brain regions into one large population, the [*D*(*R*), *D*(*C*)] correlation that is present in each area disappears (Figs. 5a2 - a3 b2 - b3, compare orange and green violin plots), consistent with what was seen in simulations (Fig. S8b, green asterisk). One potential cause of this result is the single-neuron differences (in firing rates and variances) between the two areas overshadowing the changes of activity across experimental conditions and obscuring the correlation between *D*(*R*) and *D*(*C*). To test this, we recalculated *D*(*C*) using only the variances (or equivalently, setting the off-diagonal elements of the covariance matrices to 0). In this case, the [*D*(*R*), *D*(*C*)] relation persists (Fig. 5a2 and b2, red violin plots), suggesting that the observed breakdown of our theory is at least partly due to the correlation structure in the multi-area population. This structure likely stems from the interaction between brain regions significantly violating the free independence assumption we used to derive Eq. (4) (as in Fig. 2). Similar results are also found across all pairs of areas along the mouse visual cortical hierarchy (Fig. S9; hierarchy defined according to [43]). Neighboring areas along the hierarchy exhibit especially low [*D*(*R*), *D*(*C*)] correlations in their combined activity (Fig. S9b), which may point to stronger coordinations between them yielding more structure in their covariability. Together, the above results demonstrate that our model is most readily applicable at the local circuit level, but additional considerations are needed when there are significant macroscopic structures present in the network such as connectivity differences between brain areas.

**Fig. 5:**
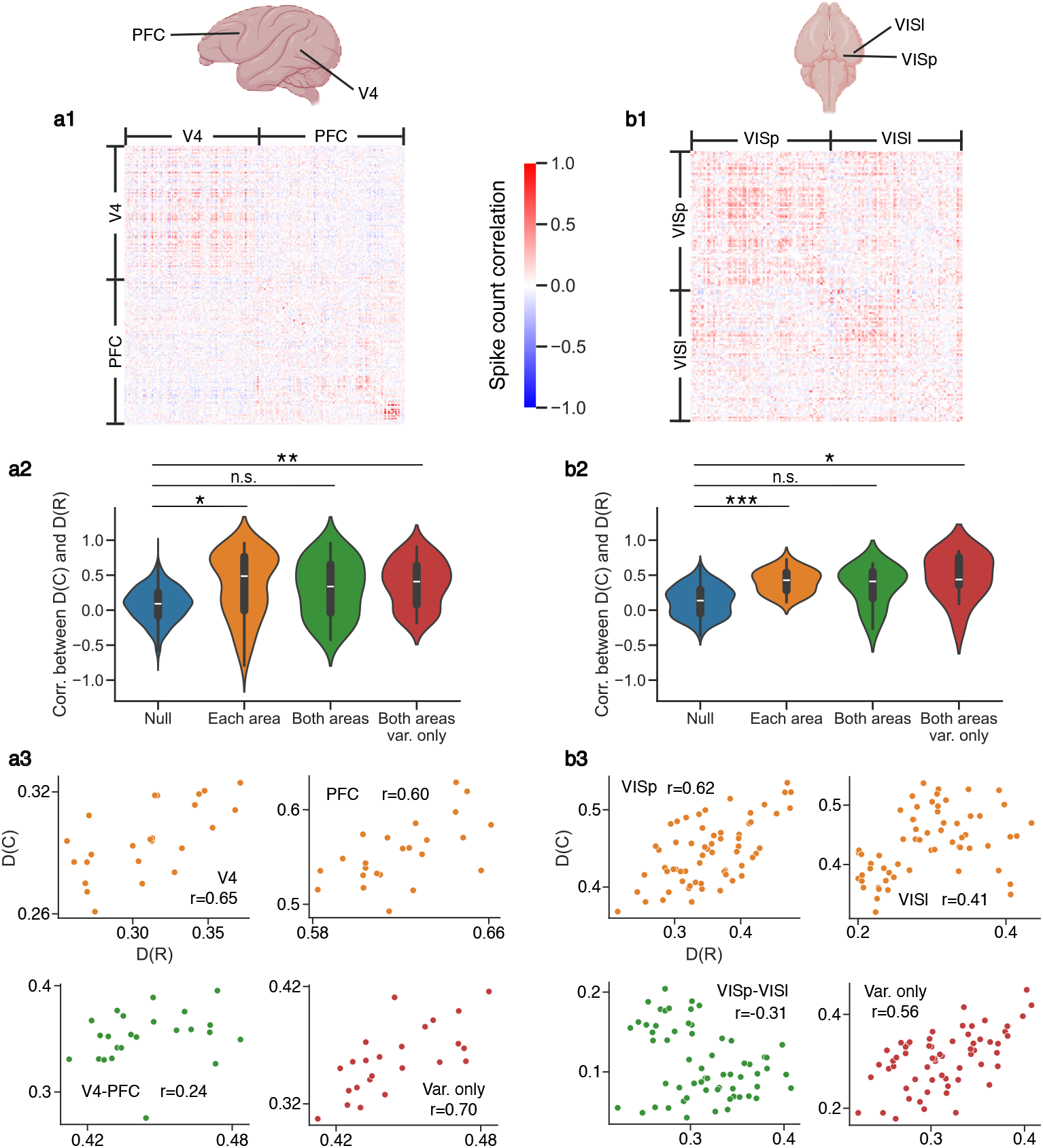
Neuronal populations that span multiple brain areas breaks the correlation between *D*(*R*) and *D*(*C*). (**a**) Data from monkey PFC and V4 [40]. (**a1**) Example correlation matrix (with diagonal elements suppressed). The correlation matrix is used over the covariance matrix for visualization purposes only. (**a2**) Distribution of the [*D*(*R*), *D*(*C*)] correlation across sessions for shuffled data, for each area separately (i.e. calculating *D*(*C*) using each of the two diagonal blocks of the full covariance matrix), for the combination of the two areas (i.e. calculating *D*(*C*) using the full covariance matrix), and for taking into account only the variances but not covariances of the two areas (i.e. calculating *D*(*C*) using the diagonal elements of the full covariance matrix). The correlation value is significantly positive for each area separately (median test, *p* = 0.011). Combining the two areas renders the [*D*(*R*), *D*(*C*)] correlation indistinguishable from the null (median test, *p* = 0.24), but not if only variances are considered (median test, *p* = 0.0032), suggesting the effect cannot be explained solely by single-neuron heterogeneity between areas. (**a3**) Example [*D*(*R*), *D*(*C*)] scatter plots from the distributions above taken from the same session. (**b**) Same as (a), but with data from mouse VISp and VISl [42]. Median tests between the three distribution pairs in (b2) yield *p*-values 1.1 × 10^−6^, 0.11, and 0.015 respectively.

### Diverse mean response works synergistically with low-dimensional variability to improve neural coding

The distribution of trial-to-trial fluctuations is critical to consider when measuring the performance of a population code for an input stimulus [8, 18, 46]. Consider the neuronal responses to two stimuli with means *R*_1_, *R*_2_ and noise covariances *C*_1_, *C*_2_. To impose stability on the network dynamics and rule out runaway activity, we consider the mean responses to have a fixed population average. This restricts *R*_1_ and *R*_2_ to lie on a simplex in the neuronal response space (Fig. 6a, yellow region). We would like to understand how mean response uniformity *D*(*R*) affects the neural code. Within the stability simplex, the mean responses with the same *D*(*R*) are defined by the intersection of the simplex with a sphere with a certain radius (Fig. 6a, magenta region), forming a codimension-2 (i.e. (*N* − 2)-dimensional) manifold *S* in neuronal response space (Fig. 6a, blue region). As *D*(*R*) gets smaller, the size of *S* gets larger so that there is more space to distribute the responses to different stimuli. At the same time, our result (Fig. 1c) suggests that the effective dimension of the population variability in both response distributions decreases, making it less likely for the trial-to-trial fluctuations to interfere with the coding dimension when the overall amount of variability is constant (compare Fig. 6b1 and b2). In sum, a diverse mean response (as measured by a small *D*(*R*)) works synergistically with low-dimensional variability (as measured by a small *D*(*C*)) to improve the population code. The synergy originates from two different aspects of neural representation with distinct contributions to decoding that are connected through network mechanisms.

**Fig. 6:**
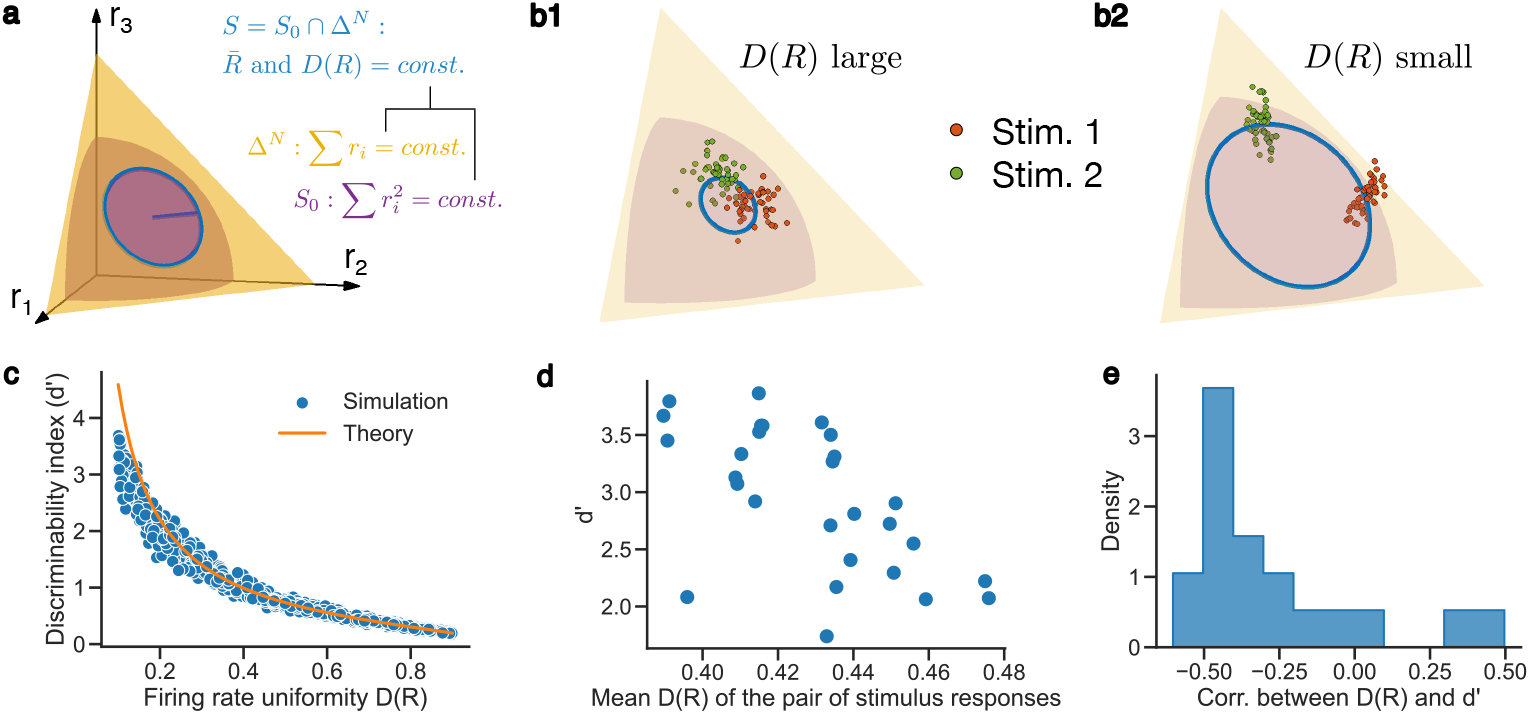
Computational implications of our result. (**a**) Schematic of the region in neuronal response space with constant firing rate uniformity *D*(*R*), which is the blue circle defined by the intersection of the sphere of constant second moment and the simplex of constant first moment. The radius of the blue circle is proportional to the standard deviation of the firing rates. (**b**) Schematic of how smaller *D*(*R*) improves the population code by enlarging the coding space (given by the size of the blue circle) and reducing the effective dimension of population variability. The red and green point clouds correspond to the neuronal response distributions to two stimuli, which the downstream decoder needs to distinguish between. (**c**) Simulation of how population codes (*N* = 100) with more diverse mean response facilitates decoding as measured by the discriminability index (d’). The theory is given in Eq. (S22). (**d**) An example session of the mean *D*(*R*) of the neuronal responses in mouse VISp [42] to two visual stimuli plotted against the estimated d’ between them. (**e**) The histogram across sessions of the correlation between the mean *D*(*R*) and d’ as in (d). It is overwhelmingly negative (*p <* 0.001).

The performance of a downstream decoder can be quantified by the (linear) discriminability index (*d*^′^). We calculated *d*^′^ as a function of *D*(*R*) when the covariance matrix of trial-to-trial fluctuations is given by Eq. (3) (Supplementary Information E). Indeed, *d*^′^ factors into two terms corresponding to the effects of mean separation and noise geometry discussed above, both of which decreases substantially as *D*(*R*) becomes larger (Eq. (S22), Fig. 6c). To verify this, we estimated the *d*^′^ of neuronal activity in mouse primary visual area (VISp) in response to pairs of drifting grating stimuli (Methods) and examined how the value of *d*^′^ for each stimulus pair relates to the average *D*(*R*) of the pair (Fig. 6d). Satisfyingly, *d*^′^ is negatively correlated with firing rate uniformity *D*(*R*) (Fig. 6e), agreeing with our prediction.

### Diverse mean response and lower-dimensional variability in states of enhanced information processing

The evidence presented above (Fig. 6e) can be viewed as describing the geometry of the neural code and could depend on which neurons are sampled in the experimental recording. To further investigate *D*(*R*)’s effect on the entire computational process, we turn to neural data spanning different behavioral states where the caliber of information processing is expected to change.

In addition to data collected during the visual categorization task (as described in Fig. 3b), which we call the “evoked state”, we also collected resting state data for an extended period of time (∼ 1 hour) when no task was given, which we refer to as the “spontaneous state”. Our theory would predict that neuronal activity has a more diverse mean and a lower-dimensional variability in the evoked state in order to better code for task related sensory stimuli, motor responses and other task variables. Indeed, there is a clear clustering of the data where both *D*(*R*) and *D*(*C*) are large for the spontaneous state and small for the evoked state (Fig. 7a, top), consistent with the prediction of our theory. The same state-dependent clustering is also present in a dataset from visual areas of mice [42] where the spontaneous state is the same as before and the mice are shown drifting grating stimuli in the evoked state (Fig. 7a, bottom). A reduction of effective dimension by stimulus presentation has also been reported in previous work [48].

**Fig. 7:**
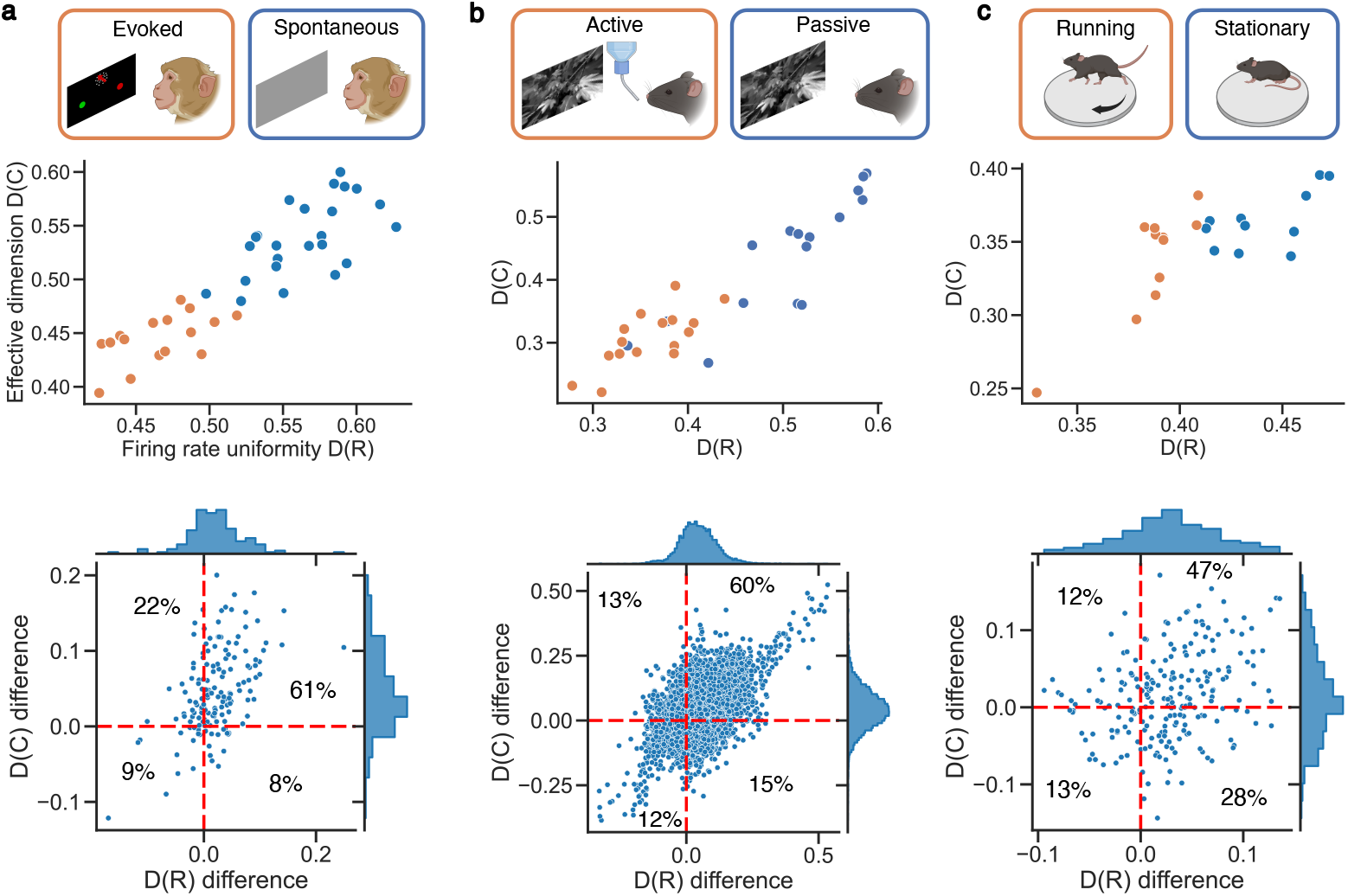
Changes of *D*(*R*) and *D*(*C*) with brain states. (**a**) Data contrasting the evoked state where the animal receives visual stimulus and the spontaneous state where the animal rests freely. The top plot was recorded in monkey FEF. The bottom plot is a summary of mouse visual area data from [42]. The areas included are VISp, VISl, VISrl, VISal, VISam, VISpm, VISmma, VISli, LP, LGd, and SGN. It shows the mean of *D*(*R*) and *D*(*C*) in the spontaneous state subtracted by those of the evoked state. (**b**) Similar to (a) but for the active state where the mice are performing a change detection task versus the passive state where the same sequence of stimuli is shown but no task is present. All data comes from [44]. The top plot is from VISal. The bottom plot shows the mean of *D*(*R*) and *D*(*C*) in the passive state subtracted by those of the active state for all recorded areas. (**c**) Similar to (a) but for the comparison between the state where the mice are running versus the state where the mice are stationary. Trials where the mean running speed of the mouse is greater than 1 cm/s are classified as “running” while others are classified as “stationary” [47]. All data is from mouse VISp in [42]. The bottom plot shows the mean of *D*(*R*) and *D*(*C*) when the mice are stationary subtracted by those when the mice are running.

In another dataset [44], the same sequence of visual stimuli was shown twice to the same mouse in two blocks, where it gets a reward for reacting to stimulus change in the first block (which we call the “active state”) but not the second block (which we call the “passive state”). In the active state, in order to perform the change detection task accurately within the given time frame, the brain needs to not only encode the sensory signal more sharply, but also be more task-engaged to quickly make a decision and transform it into motor command. Consistent with this, we see a smaller *D*(*R*) and *D*(*C*) across the brain in the active state compared to the passive state (Fig. 7b).

In mice, locomotion is associated with both better performance in visual detection and enhanced neuronal response in the visual cortex of mice [49]. Our theory would then predict more diverse mean response and lower-dimensional noise when the mouse is running to support the behavioral improvement. We indeed observe smaller *D*(*R*) for the locomotive state in mouse VISp [42] (Fig. 7c). Interestingly, *D*(*C*) is only smaller during locomotion when we remove the outlier eigenvalues, but the relation is flipped when we consider the full eigenvalue distribution (compare Fig. 7c with Fig. S10c). This is not the case for the previous two types of states we considered, where the results are consistent with or without the outliers removed (compare Fig. 7a-b with Fig. S10a-b). This suggests the presence of some low-rank modulation of population covariability associated with locomotion signals that affects the effective dimension in the opposite direction as what is caused by changes in the firing rate distribution, which is distinct from what happens for the evoked/spontaneous and active/passive states.

## Discussion

In this paper, we derived a positive relation between the diversity of mean response *D*(*R*) and the effective dimension of its population covariability *D*(*C*) (Fig. 1c), which was then tested with a wide range of data (Fig. 4). The key tool in our analysis is random matrix theory, in particular free probability theory applied to random matrices [28]. It enabled us to combine high-dimensional random variables in a nontrivial way without having to reduce the dimensionality of the system with approximation methods like mean-field theory which inevitably loses information in the process. Random matrix theory has already seen extensive applications both in neuroscience [29, 50–54] and in biology at large [55, 56]. However, most often the only random structure considered in the model is the interaction between the units in the system. Our result suggests that free probability can be very useful when trying to integrate multiple sources of randomness including complex external inputs and internal interactions.

Random matrix theory often produces results that have broad universality properties, meaning that they apply to a large class of matrix distributions satisfying some loose constraints. We developed our theory using i.i.d. Gaussian connection matrices *W* in both the feedforward and the recurrent model, but our results should hold as long as *W* is asymptotically a circular element and is freely independent from *R*. One sufficient condition is for the elements of *W* to be i.i.d. with all moments finite, and our result can be applied immediately by matching the variance. While real neuronal networks never satisfy these assumptions in the strict sense, the extent to which the [*D*(*R*), *D*(*C*)] relationship predicted by our theory holds can be an indirect indicator of the structures in the population’s connectivity and external input (or lack thereof). In the datasets we analyzed, we observe a clear positive correlation between *D*(*R*) and *D*(*C*) when neurons are localized to a single brain region (Fig. 4) but not when neurons span multiple areas (Fig. 5), reflecting the strong patterns present in interareal connections. This set of contrasting results highlight the free independence assumption as a key mathematical structure of interest in understanding population covariability in neuronal circuits.

While the participation ratio *D*(*C*) has been widely employed in neuroscience research [24, 29, 30, 57– 62], there are a wide range of other measures for the effective dimension of population activity [63, 64]. The key is to use the most appropriate definition for each scenario. At least in the context of fine discrimination tasks, we have shown that *D*(*C*) is a reasonable choice as it is directly related to the discriminability index *d*^′^ (Supplementary Information E). The relevance of *D*(*C*) is further supported by its reliable change across brain states in multiple neural datasets (Fig. 7). In particular, one should not interpret say *D*(*C*) = 0.4 in a population of 10,000 neurons as stating there being 4,000 dimensions in its activity each corresponding to some behaviorally meaningful variable. Rather, it is a measure of how concentrated the variability is around some lower-dimensional subspace, which is a statement about its geometry.

Previous works have sometimes produced seemingly different predictions on how the effective dimension depends on network diversities. Susman et al. [65] and Clark et al. [66] predict that the effective dimension of the network dynamics increases with connection weight variance (as measured by larger *g*), and Gast et al. predicts that the more heterogeneous the neurons are in terms of their firing thresholds, the larger the effective dimension [67], both of which disagree with our prediction in Eq. (4). The main difference between their models and ours is that in their networks population covariability is internally generated from network dynamics and is measured by evolving the system for some length of time. By contrast, in our model variability is fundamentally inherited from input noise and is measured by repeating the same experimental trial multiple times. In their models, the network is expanding a temporally one-dimensional input, as is typical in reservoir computing, while our model is responding to input of a certain dimensionality *D*(*C*_0_) and can only reduce the effective dimension with network heterogeneities. It is possible that population covariability in the dynamical setting (such as what would be obtained by calculating a covariance matrix across time in the production of a motor movement) may behave differently. Interestingly, we observed similar trends from spontaneous data as well (Fig. 4a, f, g, k), where we calculated the covariance matrix across time (since there was no notion of trials). This suggests that the neuronal circuits being recorded are not in a regime described by Gast et al. in the spontaneous state.

The diverse mean response discussed here is directly related to sparse coding [5], as the Treves-Rolls measure *D*(*R*) used here was originally developed to characterize the sparseness of neural codes [20–22]. There are many benefits to a sparse code [5], and our result provides another perspective on this classical idea. A natural question to ask is why neuronal responses do not always occupy the small *D*(*R*) regime, at least in sensory areas. One reason might be to keep the population code smooth with respect to stimulus parameters [68]. As *D*(*R*) gets smaller, the codimension-2 manifold *S* (Fig. 6a) becomes topologically more complex. In particular, it acquires “holes” and becomes “less and less connected” in the sense of homotopical connectivity, eventually becoming disconnected (Eq. (S21)). Representation of stimulus features amounts to embedding the feature space into the neuronal response space. A smoother embedding may inevitably need to include regions of larger *D*(*R*), and the full neural manifold that encodes all stimuli of interest likely spans a range of *D*(*R*) values. How different stimuli are mapped to codes with different levels of sparseness is an interesting question. Our results here would predict that stimulus sets that require higher precision in discerning the finer details would correspond to sparser codes. This prediction is supported by a recent study in which training mice to perform a task sparsified the population responses to task-relevant orientations to a much greater extent than non-task orientations [69]. More generally, sparsification of neuronal response due to active learning or repeated passive exposure to sensory stimuli has been observed in the visual cortex [22, 69, 70], somatosensory cortex [71], olfactory bulb [72], auditory cortex [73] and hippocampus [74], and has also been produced in a model with synaptic plasticity rules inferred from data [75]. Thus, a sparsification of neuronal response may be reserved for computations that require a fine discrimination of stimulus features.

In our model *W* and *C*_0_ are independent of *R*, so the coding dimensions are independent of the directions of variability in neuronal response space. In this setting, the effective dimension is a very good summary of the geometry of neuronal variability. However, it is possible that the modes with the largest eigenvalues are the most important for coding, and the bulk portion of the eigenvalue distribution studied here is not as critical. However, the largest eigenmodes of primary visual cortex activity have been shown to be orthogonal to coding dimensions in both monkeys and mice [76, 77], suggesting that they do not interfere with visual scene representation (but see [78]). We would further argue that the bulk portion of the eigenspectrum is almost always important. These eigenmodes can be interpreted as originating from the component of the recurrent connectivity that is unrelated to the task at hand but may serve other functions in other contexts, and is high-rank in nature. Low-rank noise can be eliminated downstream if it is shared across areas [79] with adjustments in only a few subspaces, but the bulk of the noise, even with a magnitude not as large, cannot be easily removed due to it being distributed across many dimensions. Reducing the effective dimension by either increasing the diversity of the mean response or relying on the heterogeneity of intrinsic neuronal properties [80, 81] then presents a global strategy of improving the population code without any fine-tuning in the connectivity. To be sure, the correlation between *W*, *C*_0_ and *R* definitely has an enormous impact on the neural code, as has been extensively studied [17, 82], and it would be interesting to investigate how to incorporate this aspect into our framework.

Much attention has been devoted to understanding how the structure of population covariability and its effective dimension *D*(*C*) in particular can be flexibly controlled in the brain [19]. In our model, gain modulation (i.e a change in *L*) is the root cause of how *C* changes across brain states, as plasticity is assumed to operate on a slower timescale. This can be achieved by changing the mean input vector to the circuit, either feedforward or feedback, which in turn changes the operating point of the system. This amounts to changing the “effective connectivity” of the network, whose effect on neuronal variability has also been studied in previous works [16, 26, 83]. The dominant factor behind a change in effective dimension may be different depending on the brain area and the task setting, and needs to be examined on a case-by-case basis. The mechanism proposed by our model is perhaps the easiest to check, since *D*(*R*) can be immediately calculated with the same data that was used to estimate *D*(*C*) without any additional procedure. Our result thus sets a basic expectation for changes in *D*(*C*) based on changes in *D*(*R*). When the effective dimension *D*(*C*) changes across conditions or states, one can first check if the changes in *D*(*R*) accounts for it (as we show in Figs. 7 and S10). If not, one can then look closer at the structure of the covariance matrix *C* (such as large outlying eigenvalues which could correspond to low-rank feedback modulation [84]) or perform further experiments to reveal other factors responsible for controlling the effective dimension of the variability.

In summary, we derived in feedforward and recurrent circuit models that the more diverse the mean response is in the network, the lower the effective dimension of population covariability. We verified this result in neural data from multiple brain areas from monkeys, mice, and human participants. We showed that a diverse mean response works synergistically with low-dimensional noise to improve the population code, which is again supported by experimental evidence. Our work provides a mechanistic framework for understanding how different sources of heterogeneity in a network interact to determine the geometry of neuronal variability.

## Methods

### Non-human primate experiments

#### Subjects and behavioral tasks

Two male rhesus monkeys (Macaca mulatta) participated in this experiment: Monkey M, 21-24 years old, ∼ 10 kg; Monkey S, 9-12 years old, ∼ 13 kg. Both monkeys were implanted with a headpost and trained to perform the memory-guided saccade (MGS) task and the rapid categorization (RC) task (described in detail in the next paragraph). For all tasks, the monkeys were head restrained and seated in a primate chair inside an isolation box (Crist Instruments) or in a dedicated experimental room. Visual stimuli were displayed on a 21-inch color CRT monitor (1280 × 1024 resolution, 75 Hz refresh rate, 57cm viewing distance). Gaze positions were tracked using an Eyelink 1000 optical eye tracker (SR Research, Ottawa, Canada) at a sample rate of 1 kHz. Task events, stimulus presentation, behavioral signals, and reward delivery were controlled by the MATLAB-based toolbox NIMH MonkeyLogic [85, 86] on a Windows 10 machine. After training, both monkeys were implanted with two custom titanium recording chambers in the right hemisphere (Rogue Research): one chamber was positioned over the frontal eye fields (FEF) and the other was positioned over both the lateral intraparietal area (LIP) and the superior colliculus (SC) based on stereotaxic coordinates from pre-implantation MRI scans. All surgical and experimental procedures adhered to the guidelines of the University of Chicago’s Institutional Animal Care and Use Committee and the National Institutes of Health.

#### Memory-guided saccade task

To identify visual and motor receptive fields of LIP, FEF, and SC, we used the MGS task [87–89]. Monkeys initiated trials by acquiring and maintaining fixation on a central white fixation point (0.2° radius) within a 2.5° fixation window throughout the trial. After 500 ms of fixation, a 0.6° white square target was briefly presented at one of eight peripheral locations (7° eccentricity) for 300 ms. After a 1000 ms delay period, the fixation cue disappeared, and monkeys made a saccade to the remembered location of the target for a juice reward delivered through a custom-made juice-spout. Recording coordinates in all areas were selected if they contained neurons with visual or motor receptive fields to the left of the fixation point (contralateral to the recorded hemisphere).

#### Rapid categorization task

Similar to the one-interval categorization task reported in previous experiments [89], monkeys categorized random-dot motion stimuli into one of two categories with a saccadic eye movement. The motion stimuli consisted of 5° diameter circular patches of ∼ 100 high-contrast dots moving at 12°/s with 100% coherence. A set of six directions spanning 360° range of motion was arbitrarily divided into two categories, and monkeys must report the learned motion category with a saccade to a red or green color target (Red: 75°, 135°, 195°; Green: 15°, 255°, 315°). Monkeys initiated trials by acquiring and maintaining fixation on a central fixation point (0.2° radius) within a 2.5° fixation window throughout the trial. After 500 ms of fixation, a place-holder random dot motion stimulus with 0% coherence (5° diameter) was positioned 7° to the left of the fixation point. This place-holder stimulus is used to avoid the bottom-up attention capture of the full coherence motion stimulus and any transients of neuronal activity corresponding to the appearance of any stimulus in the receptive field of the neuron. After 300 ms of continued fixation, red and green targets (1° square) appeared at positions 7° above and below the fixation point. Importantly, the relative positions of the color targets were randomized on every trial to decouple specific saccade directions from motion categories (e.g. the red target can either be above or below the fixation point on different trials). After 350 ms of continued fixation, the 0% coherence noise stimulus is replaced by a 100% coherence motion stimulus and simultaneously, the fixation point disappears, signaling to the monkeys that they must make a saccade within 500 ms. If they saccade to and fixate on the correct color target for 300 ms, they receive a juice reward delivered through a juice spout.

#### Passive viewing (spontaneous state)

To measure the spontaneous patterns of activity in each area, recordings continued after performing the previous tasks while monkeys passively sat in a dark environment for 30-60 minutes without any visual stimuli on the monitor, sounds, or behavioral requirements. Gaze positions were continuously tracked to monitor for spontaneous eye movements or for periods of sleep.

#### Electrophysiological recordings

Electrophysiological procedures are similar to previous studies from our group [89]. Recordings were conducted in separate sessions using combinations of 24-channel or 32-channel linear arrays (Plexon V-probes and Plexus S-probes) and a NAN microdrive system (NAN Instruments). Up to six probes were used to simultaneously record LIP, FEF, and SC (two per area, up to four in one recording chamber). Neurophysiological signals were amplified, digitized, and stored for offline spike sorting (Plexon) using the MATLAB-based Kilosort 2 toolbox, which ran on a Windows 10 PC with a NVIDIA RTX 3070 graphics card [90]. Putative neuron clusters identified via Kilosort 2 were curated using a Python toolbox Phy [91].

### Human experiment

#### Participants

This study was conducted under an Investigational Device Exemption from the U.S. Food and Drug Administration and approved by the Institutional Review Boards at the Universities of Pittsburgh and Chicago. The clinical trial is registered at ClinicalTrials.gov (NCT01894802). Informed consent was obtained before any study procedures were conducted. Participant C1 (male, m), 55-60 years old at the time of implant, presented with a C4-level ASIA D spinal cord injury (SCI) that occurred 35 years prior to implant. C1 retains partial function of the contralateral arm and risk but can only achieve hand movements with tendonesis grip. Participant C2 (male), 60-65 years old at time of implant, presented with a C4-level ASIA D spinal cord injury (SCI), along with a right brachial plexopathy, that occurred 4 years prior. C2 retains some shoulder function and wrist function.

#### Cortical implants

We implanted four microelectrode arrays (Blackrock Neurotech, Salt Lake City, UT, USA) in each participant. The two arrays (one medial and one lateral array) in Brodmann’s area 1 of S1 were 2.4 mm x 4 mm with sixty 1.5 mm long electrode shanks wired in a checkerboard pattern such that 32 electrodes could be recorded from or stimulated. The two arrays in motor cortex were 4 mm x 4 mm with one-hundred 1.5 mm long electrode shanks wired such that 96 electrodes could be used to monitor neuronal activity. The inactive shanks were located at the corners of these arrays. Two percutaneous connectors, each connected to one sensory array and one motor array, were fixed to the participant’s skull. We targeted array placement during surgery based on functional neuroimaging (fMRI) or magnetoencephalography (MEG) of the participants attempting to make movements of the hand and arm within the constraints of anatomical features such as blood vessels and cortical topography – see [92] for a full description of the implantation approach.

#### Center-out reaching task

To facilitate arm movements given the reduced motor function of each participant, each participant’s arm was rest upon a forearm support with ball bearings to reduce friction and the need to lift the arm during movement. Participant C1 was provided 8 target positions equally spaced around a circle 30 cm in diameter. Due to reduced shoulder function of participant C2, 4 targets were sub-selected (45, 90, 135, 225 degrees). Each trial was composed of 5 phases, all 1 second in duration and indicated by auditory tones. The ‘cue phase’ was indicated with a long auditory tone at which point the trial target would be indicated to the participant via text on a TV (e.g. “top-left”, “bottom-right”). After 1 second the ‘movement phase’ began as indicated by a short tone at which point each participant would then slowly move their arm to the indicated target over the 1 second phase period such that they arrived at the target when the tone for the next phase occurred. Participants then remained stationary during the ‘wait phase’ until the ‘return phase’ tone played at which point they slowly returned to the center position. Finally, the participants waited for another second before the next trial began. Participants were allowed to pause as requested. Each participant completed 11 blocks of reaches in which each trial was repeated 10 times. Consequently, participant C1 completed 880 reaches and participant C2 completed 440 reaches.

#### Passive ‘task’

To contrast with the active nature of the center-out reaching task we also had participants relax and watch TV (a nature documentary) for approximately 60-90 minutes. Participant C1 remained engaged with the documentary for the entire duration and often made comments to the experimenters. Participant C2 was less engaged and fell asleep approximately half way through the documentary.

#### Data acquisition and processing

Cereplex ES headstages (Blackrock Neurotech, UT, USA) were used to connect each pedestal to a NeuroPort system (Blackrock Neurotech, UT, USA). Voltages were sampled at 30 kHz and high-pass filtered (750 Hz) with a 1st order Chebychev filter. An RMS threshold of -4.5 was used to detect spikes from the continuous signal. Post-hoc offline sorting of single units was performed using Offline Sorter (v4.7.1, Plexon, TX, USA).

### Measure of mean response uniformity and effective dimension

The Treves-Rolls measure [20–22] for quantifying the uniformity of firing rate in a neuronal population is defined as

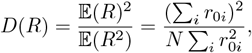

where *r*_0*i*_ is the firing rate of neuron *i*. When all neurons have the same firing rate, we have *D*(*R*) = 1, marking the most uniform activity level. As neurons fire at more and more diverse rates, *D*(*R*) becomes lower. This can be seen as an interpolation of the cases where *M* neurons fire at the same rate while other neurons are silent, which has *D*(*R*) = *M/N*.

The participation ratio [23, 24] as a measure of the effective dimension of population covariability is defined as

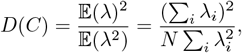

where *λ*_*i*_ is the *i*’th eigenvalue of the spike count covariance matrix *C*. When variability is isotropic in neuronal response pace with the same magnitude in all directions, we have *D*(*C*) = 1, signifying the maximum effective dimension. As the noise is distributed more and more unevenly in different directions, *D*(*C*) becomes lower. This can be seen as an interpolation of the cases where *M* eigenvalues of *C* have the same positive value while the other eigenvalues are 0, i.e. where the variability is restricted to an *M* -dimensional subspace, which has *D*(*R*) = *M/N*.

The formal similarity of these measures is not a coincidence, as the participation ratio is exactly measuring the uniformity of noise across orthogonal dimensions.

### Simulating firing rates with prescribed *D*(*R*)

Here we describe how to sample firing rates from a log-normal distribution ℒ𝒩(*µ, σ*^2^) with a given mean 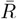 and uniformity *D*(*R*) as we did in Fig. 3a). For *X* ∼ ℒ𝒩(*µ, σ*^2^), we have

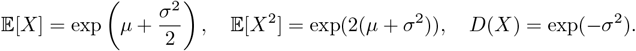

Then

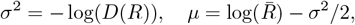

gives us the desired parameters for the log-normal distribution.

### Simulations with spatially structured connectivity and input noise

To make Fig. 2, we perform similar procedures as Fig. 3a except *W* and *C*_0_ are generated in a spatially dependent manner. Specifically, each neuron in the population is randomly assigned a location in a two-dimensional grid of length 1. For neurons *i* and *j* that are distance *d*_*ij*_ apart, we let the connection between them be *w*_*ij*_ = *w*(*d*_*ij*_) and the input covariance between them be 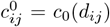, where *w* is the Laplacian of a Gaussian with mean 0 and standard deviation *α*_*W*_, and *c*_0_ is a Gaussian with mean 0 and standard deviation 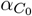. The reason we use the Laplacian of a Gaussian for *w* is to keep the mean of the elements of *W* near 0. In this setup, the region in the upper left corner of Fig. 2b with *α*_*W*_ close to 1 and 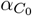 close to 0 (exemplified by Fig. 2c) is where *W* and *C*_0_ are closest to being rotational invariant and where the [*D*(*R*), *D*(*C*)] correlation predicted by our theory is expected to hold. The *W* and *C*_0_ generated this way are Euclidean random matrices [93]. Note that we are not explicitly modeling the covariance matrix *C* itself as an Euclidean random matrix, as was done in previous work [52].

### Data processing

We describe here the general approach of processing the population electrophysiology data shown in Fig. 4. When there is a trial structure present during the recording, such as the presentation of stimuli or the execution of some motor action, we first separate the trials into groups with the same experimental condition. This includes the same sensory stimuli, task cue configuration, behavior of the subject etc. Within each group, we further partition the trials into different epochs delimiting different stages of a trial such as fixation, stimulus presentation, and behavioral response. Finally, we find a 100 ms bin in each epoch where neuronal activity is the most stationary and count the number of spikes in the bin for each neuron. Then we obtain a set of spike count matrices of size (number of trials × number of neurons) for each experimental condition and trial epoch. Each of these matrices are regarded as samples from some distribution of neuronal response, from which we can calculate the firing rate distribution and the covariance matrix across trials for this operating point of the system. This in turn gives us *D*(*R*) and *D*(*C*). For example, 4 conditions with 5 epochs in each trial would give us 20 points in the [*D*(*R*), *D*(*C*)] scatter plot. For resting state data where the subject is spontaneously behaving without any trial structure, we take the system to roughly be at the same operating point in each minute-long recording, and use the slow drift of the system’s state across time [94] to sample different [*D*(*R*), *D*(*C*)] pairs in place of experimental conditions and trial epochs. That is, we calculate the mean responses and covariances for each minute in the recording by binning it into 100 ms spike counts. Then for example, a 30-minute long spontaneous state recording would produce 30 points in the [*D*(*R*), *D*(*C*)] scatter plot. An additional point to note is, previous work has shown in monkeys that whether the eyes are open or closed can have significant impact on the spiking activity of neurons including its effective dimension [95, 96]. We therefore removed the portion of the data where the monkey has its eyes closed or is making a saccade using eye-tracking data.

#### Fitting the eigenvalue distribution to data and detecting the outliers

As discussed in the main text, our model only accounts for the bulk of the covariance matrix eigenvalue distribution and we need to remove the outlier eigenvalues before verifying our theory. We do this by fitting our model to the data. Given firing rate distributions *R*_*i*_ where *i* indexes different operating points (due to different experimental conditions and trial epochs as discussed earlier) and covariance matrix eigenvalues *λ*_0*i*_ ∈ ℝ^*M*^ from data recorded from a population of *M* neurons, we need to fit two parameters: the connection diversity parameter *g* and the number of neurons in the underlying population *N* . Synaptic connectivity is assumed to stay constant in each recording session, so *g* and *N* do not depend on the operating point *i*. Since we are considering spatial sampling here (i.e. *M* < *N*), there is no analytic solution to the eigenvalue problem (Supplementary Information D.2), so we need to use a search algorithm to find the combination of *g* and *N* that best explains the data. We use Bayesian optimization [97] to perform the search. The cost function is defined by

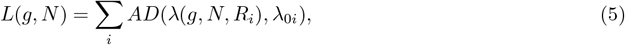

where *AD* denotes the Anderson-Darling statistic [98] which measures how close two distributions are to each other and *λ*(*g, N, R*_*i*_) is the samples generated by first simulating an *N* × *N* covariance matrix with parameter *g* according to Eq. (3) (with *p* = 2 and *C*_0_ = *I*) using the firing rate distribution *R*_*i*_ and then randomly picking a *M* × *M* principle submatrix to calculate eigenvalues. This procedure is done many times (typically ∼ 100*M*) to ensure a good sampling of the model distribution. Each repetition has either a different realization of connectivity *W* or a different sampled submatrix. There is an unknown scaling parameter as summarized by *k*_0_ in Eq. (S6). We eliminate its effect by matching the means of *λ*(*g, N, R*_*i*_) and *λ*_0*i*_ before calculating the Anderson-Darling statistics. Note that this simulation process is inherently stochastic since the generated eigenvalue samples depend on the realizations of the weight matrix *W* and which *M* neurons are sampled from the total size-*N* population (even after repeating the simulation for many times), but this stochasticity is allowed by the Bayesian optimization algorithm.

There is sometimes degeneracy between *g* and *N* in that there may be a continuous set of combinations that produce similarly small cost functions. For example, Fig. S4c shows a wide distribution of (*g, N*) combinations that are close to optimal in an example session in monkey LIP. This degeneracy is likely responsible for the wide distribution of fitted *ĝ* values across sessions for each brain area, to the extent that most areas have a *ĝ* distribution indistinguishable from the aggregate over all areas (Fig. S4d). However, for the purpose of getting a null model for the eigenvalue distribution and identifying the outliers in the data, any good pair of *ĝ* and 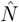 can be used to generate 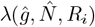 that sufficiently captures the bulk of *λ*_0*i*_. Once we have a good sample of *λ*(*g, N, R*_*i*_), we can identify the eigenvalues in the neural data that lie outside of its support as outliers. Alternatively, when *M* is relatively small, even a data eigenvalue within the null model support may be unexpected if it is very large. Let *λ*_1_ be a large eigenvalue in neural data. We can set a reference *p*-value *p*_0_ and calculate the likelihood of obtaining a sample from 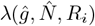 no smaller than *λ*_1_ in *M* draws. If that likelihood is smaller than *p*_0_, we say *λ*_1_ is an outlier. In our study we adopt the second method with *p*_0_ = 0.01. We typically find 0-2 outliers in data. All of the *D*(*C*) values presented in the figures are calculated after removing these outlier eigenvalues except in Fig. 4e, j, in the small boxes in Fig. 7, and in Fig. S6.

In Fig. S7c-d, we compared the fit of our model (“diverse gain”) to those of two other models. For the “uniform gain” model [29], we set *R* to be uniform across neurons when generating *λ*(*g, N, R*_*i*_) and performed the same optimization procedure as above. In particular, the number of fitted parameters is the same, and the only difference is the “uniform gain” model does not incorporate information from the firing rate distribution. For the “uncoupled” model we simply multiplied the firing rate distribution by a scalar to match its mean with the eigenvalues to get the eigenvalues that would be produced by a population of uncoupled Poisson neurons.

#### Correcting for temporal sampling

In Supplementary Information D.1, we discuss how a limited number of temporal samples/trials affects the estimation of the effective dimension *D*(*C*). To correct for this effect, we use the simple formula

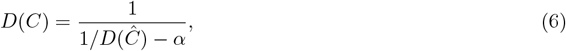

where *D*(*Ĉ*) is the naive sample covariance matrix and *α* = *N/T* with *T* being the number of temporal samples. This recovers the effective dimension of the exact covariance matrix. However, this procedure only corrects for the bias due to temporal sampling, not the variance. If *T* is too small, the estimate of *D*(*C*) would still be very noisy even after the correction. In principle, the estimate of *D*(*R*) may also be noisy given limited data, although presumably to a lesser extent. To investigate the reliability of *D*(*R*) and *D*(*C*) when the number of trials is limited, we use a bootstrapping analysis on the mouse dataset in [42, 43]. Specifically, we pick sessions and brain areas with sufficient numbers of trials, subsample the trials multiple times, and compare the result to that from the full data to characterize the bias and variability in the estimate as a function of temporal sampling. We find that *D*(*R*) and *D*(*C*) can be estimated reasonably well as long as the number of temporal samples is more than about 3 times that of the number of neurons (Fig. S4e), which is true for most of the datasets we use in this paper.

#### Spike count binning considerations

In our analysis we always pick the period of each trial when the population activity is the most statistically stationary, since it is assumed in our theoretical derivations. However, in reality neuronal response is never perfectly stationary in time, and it is unclear how critical our result depends on the choice of time window location and bin size for spike counts. To study this, we reanalyzed the monkey V4 dataset in [40] by varying which time period is used in each trial to count the spikes (Fig. S4f). In particular, the population activity exhibits a transient in the early period of the trial, making it strongly nonstationary. We find that the [*D*(*R*), *D*(*C*)] correlation does not depend significantly on either the choice of time window location or bin size (Fig. S4g, h). This demonstrates the robustness of our result to detailed data processing choices.

#### Shuffled data

We describe here how we generated the null distribution in Fig. 4d, i. What is measured by the correlation between *D*(*C*) and *D*(*R*) is how these two quantities covary when the operating point of the system changes. However, due to finite-size effect and the discrete nature of spike count data, we may observe spurious correlation between them even when we repeatedly sample spike count data from the same operating point. To control for this, we need to compare with shuffled data. Suppose we have *K* operating points in the data (i.e. *K* points in the [*D*(*R*), *D*(*C*)] scatter plot). We take the operating point with *D*(*C*) and *D*(*R*) closest to the mean of all [*D*(*R*), *D*(*C*)] pairs and generate *K* spike count matrices each having the same number of temporal samples as the data according to the firing rate vector and covariance matrix of this operating point using the algorithm in [99]. This in turn gives us *K* points in the [*D*(*R*), *D*(*C*)] plane which can be used to calculate a correlation. This is done 10 times for each brain area and recording session. The resulting set of correlation values compose the null histograms.

### Calculation of the discriminability index

For two stimulus responses with means *r*_1_, *r*_2_ and covariance matrices *C*_1_, *C*_2_, the discriminability index *d*^′^ is [100]

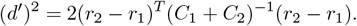

For the simulation in Fig. 6c, we first drew *r*_1_ from some log-normal distribution with a certain mean and *D*(*R*) as we did in Fig. 3a. We then sample *r*_2_ in the same manifold *S* (Fig. 6a at a distance *d*_0_(1*/D*(*R*) − 1) from *r*_1_ in a random direction. Here *d*_0_ is a scaling constant and the distance gets larger at the same pace as the overall size of *S* (Supplementary Information E). This is achieved by first sampling a random unit vector in the neuronal response space, then projecting it to the linear subspace orthogonal to both the vector of ones (to ensure it stays in the simplex Δ^*N*^) and the vector *r*_1_ (to ensure it roughly stays in the sphere *S*_0_), and finally scaling it by *d*_0_(1*/D*(*R*) − 1) to obtain *r*_2_ − *r*_1_. *C*_1_ and *C*_2_ are again generated according to Eq. (3) with *p* = 2 and *C*_0_ = *I*. Then it is straightforward to calculate *d*^′^.

To estimate *d*^′^ in neural data in Fig. 6d-e, we fit linear classifiers (logistic regression) to each pair of stimulus responses and get the cross-validated classification accuracy *a. d*^*′*^ is related to *a* by [100]

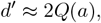

where *Q* is the quantile function of the standard normal.

## Data availability

Will be available for public use upon manuscript publication.

## Code availability

All modelling and analysis codes are provided at https://github.com/brain-math/fr-diversity-lowers-cov-dimensionality

## Acknowledgments

We thank Michael Boninger, MD, for serving as the sponsor-investigator of the BCI clinical trial conducted under an FDA Investigational Device Exemption. C.G. and J.D. are supported by the National Institute of Neurological Disorders and Stroke of the National Institutes of Health (NIH) under award numbers UH3 NS107714, R35 NS122333, and R01 NS130302. D.J.F is supported by NIH grant R01EY019041 and a Vannevar Bush Faculty Fellowship (N000141912001). O.Z. is supported by NIH grants F30EY033648 and T32GM007281. B.D. is supported by the NIH grants 1U19NS107613-01, R01NS133598 and CRCNS-R01EY034723, a Vannevar Bush faculty fellowship (N000141812002), and the Simons Foundation Collaboration on the Global Brain. This research benefited from Physics Frontier Center for Living Systems funded by the National Science Foundation (PHY-2317138). Financial support was provided via the National Institute from Mathematics and Theory in Biology (Simons Foundation award MP-TMPS-00005320 and NSF award DMS-2235451). We thank Matthew A. Smith and Shenghao Wu for help on the monkey V4-PFC dataset. Cartoon schematics created with BioRender.

## Author contributions

G.J.T. and B.D. conceived of the project, theory, and data analysis. G.J.T. performed the analytic calculations and the data analysis. O.Z., V.S. and D.J.F. performed the monkey electrophysiology recordings (areas LIP, FEF, and SC). C.M.G. and J.E.D. performed the human participant electrophysiology recordings. All authors were involved in the writing of the manuscript.

## Supplementary Information

### A The mean response and the covariance matrix are related through nonlinear transfer

Consider a population of *N* neurons. Let *x*_*i*_ be the synaptic input to neuron *i* and *r*_*i*_ be its output activity (*i* = 1, …, *N*). Denote the means and covariance matrices of *x* ∈ ℝ^*N*^ and *r* ∈ ℝ^*N*^ by 𝔼[*x*] = *x*_0_, 𝕍[*x*]= *C*_*x*_, 𝔼[*r*] = *r*_0_, 𝕍[*r*] = *C* respectively (Fig. 1a). The input and output of a neuron is related by its transfer function *f*, i.e., *r*_*i*_ = *f* (*x*_*i*_). The gain of the neuron describes the sensitivity of its output against small input changes and can be defined by linearizing around *x*_0_ as *l*_*i*_ = *f* ^*′*^(*x*_0*i*_). Then by linear approximation we have *C* = *LC*_*x*_*L*^*T*^, where *L* = diag(*l*). If *f* is nonlinear, then *l*_*i*_ would depend on *x*_0*i*_ and thus *r*_0*i*_ = *f* (*x*_0*i*_) (technically we only have an approximate relation *r*_0*i*_ ≈ *f* (*x*_0*i*_), but this is sufficient for our purposes). In turn, the covariance matrix *C* of neuronal output should depend on the mean response *r*_0_.

To be concrete, and consistent with experimental results, we use the threshold-power law as the transfer function [25]:

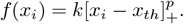

This produces the following gain matrix:

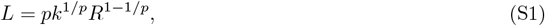

where *R* = diag(*r*_0_). Throughout the paper, we also use *R* to denote the distribution of firing rates, which is exactly the eigenvalue distribution of *R* as a diagonal matrix. Following previous work, we set the power *p* to be 2 unless otherwise stated [26]. In this case we have *L*^2^ ∝ *R*.

The specific relation between *R* and *C* (and thus *D*(*R*) and *D*(*C*)) depends on the structure of *C*_*x*_. In the simplest case of i.i.d. input noise across neurons, we have *C*_*x*_ = *I* and *C* ∝ *R*, which implies *D*(*C*) = *D*(*R*). However, whether this relation holds for other input covariances remains to be investigated. While there are in principle many ways of modeling *C*_*x*_, we take a mechanistic approach and in the following sections consider the cases of feedforward and recurrent circuits (Fig. 1c).

### B The feedforward model

#### B.1 Expression for the covariance matrix

We consider the structure of the input covariance matrix *C*_*x*_ as originating from the feedforward projection *W* ∈ ℝ^*N ×M*^ from an upstream population of *M* neurons having i.i.d. unit-variance variability (i.e., with the identity matrix *I* as the covariance matrix). Then *C*_*x*_ = *WW*^*T*^ and

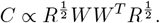

For concreteness, let us assume that the elements of *W* are i.i.d. with distribution *w*_*ij*_ ∼ 𝒩 (0, *g*^2^*/M*) for some *g >* 0 and that *W* is independent of *R*. Note that different values of *g* only scales *C* by a number and do not affect *D*(*C*) as it is a normalized quantity.

#### B.2 The covariance eigenspectrum

We use techniques from free probability to derive the eigenvalue distribution of *C* [28]. Following standard notations, we define for any random matrix *A* of size *N*

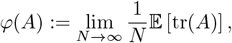

where tr(*A*) = Σ_*i*_ *A*_*ii*_ is the trace. Note that all derivations here are in the large *N* limit, and we shall not make a clear distinction between a random matrix of finite size with the object it converges to, nor shall we discuss different notions of convergence such as weak and a.s. convergence. We observe in simulations that all of our results converge for *N* at a few hundred, which is comparable to the size of the neural data we analyze.

Let *a, b*, and *c* be the corresponding variables in the non-commutative probability space (𝒞, *φ*) of *R, C*_*x*_, and *C* respectively. Note that the nonzero eigenvalues of *C* are the same as those of *C*_*x*_*R*, so *c* = *ab*. Since *C*_*x*_ and *R* are assumed to be independent and *C*_*x*_ is rotationally invariant, the product *C*_*x*_*R* is free [28]. Then their *S*-transforms satisfy

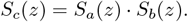

The eigenvalue distribution of *C*_*x*_ = *WW*^*T*^ follows the Marchenko-Pastur law, so *b* is a free Poisson element whose *S*-transform is known to be

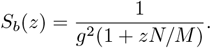

For any variable *x*, its *S*-transform *S*_*x*_(*z*) satisfies

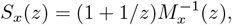

where *M*_*x*_ is the moment-generating function

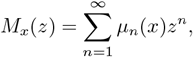

where *µ*_*n*_(*x*) is the *n*’th moment of *x*. We shall denote the *n*’th cumulant by *κ*_*n*_(*x*), and 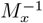 is the inverse function of *M*_*x*_. Also, we have the relation

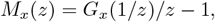

where *G*_*x*_ is the Cauchy transform defined to be *G*_*x*_(*z*) = *φ*((*z* − *x*)^−1^). The Cauchy transform is related to the eigenvalue distribution *p*_*x*_ by

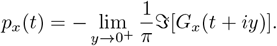

Using the above relations we have

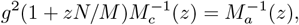

and writing *z* = *M*_*c*_(1*/z*_1_) = *M*_*a*_(1*/z*_2_) gives us the system of equations

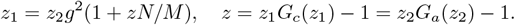

To get the probability density function (pdf) *p*_*c*_ of *c*, we need to solve for *G*_*c*_(*z*_1_) for each *z*_1_ approaching the real axis. Since *G*_*a*_ can be calculated numerically from the distribution of *R*, we can first solve

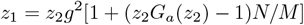

for *z*_2_ and then plug it back to get *z* and thus *G*_*c*_(*z*_1_). This is verified in simulations (Fig. S1a).

#### B.3 The effective dimension *D*(*C*)

Since *C*_*x*_ and *R* are assumed to be free, we can calculate

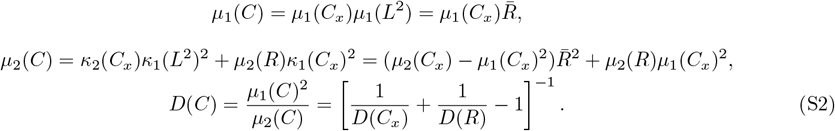

Here 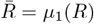 is the population average firing rate. We see that *D*(*C*) is positively related to both *D*(*R*) and *D*(*C*_*x*_). In particular, for *C*_*x*_ = *WW*^*T*^ we have

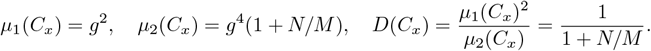

Inserting the expression for *D*(*C*_*x*_) in Eq. (S2) gives us Eq. (2).

### C The recurrent model

A key assumption in the previous section is that *C*_*x*_ and *R* are independent. This is generally not true, especially in a recurrent network where the input to each neuron is affected by the outputs of other neurons in the population. We examine this case in this section.

#### C.1 Expression for the covariance matrix

Consider a recurrent network of *N* neurons

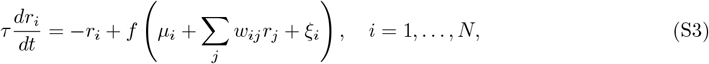

where *r*_*i*_ is the response of neuron *i, τ* is the neuronal time constant, *f* is the input-output transfer function, *µ*_*i*_ is the mean external input, *w*_*ij*_ is the connection weight from neuron *j* to neuron *i*, and *ξ*_*i*_ is trial-to-trial input noise. In vectorized notation we can write

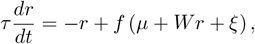

where *r* := (*r*_*i*_)_*i*_, *µ* := (*µ*_*i*_)_*i*_, *W* := (*w*_*ij*_)_*i,j*_, *ξ* := (*ξ*_*i*_)_*i*_, and *f* is applied element-wise. Note that *x* = *µ*+*Wr*+*ξ*. In each trial, assume the network evolves to a steady state

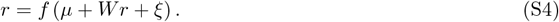

Here *r* is a function of *ξ*, which is a random variable. When the noise *ξ* is small, we can linearize to get the approximation

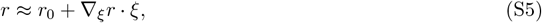

where *r*_0_ is the steady state when there is no noise, i.e.

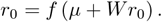

Differentiating Eq. (S4) gives us

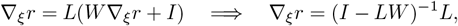

where *L* := diag(*l*_1_, …, *l*_*N*_) is the diagonal matrix of neuronal gains

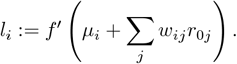

Then

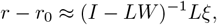

and the covariance matrix *C* is

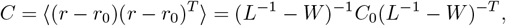

where *C*_0_ = ⟨*ξξ*^*T*^⟩ is the covariance matrix of the input noise, which we assume to be *C*_0_ = *I* in the following. This is exactly Eq. (3). We remark that a wide range of linear or linearizable systems can formally lead to Eq. (3) [29, 32].

#### C.2 The covariance eigenspectrum

We next derive the eigenvalue distribution of the covariance matrix *C* under the assumptions 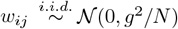, that *f* follows Eq. (1), and that *L* and *W* are independent. We again have Eq. (S1). Since we are ultimately interested in the effective dimension *D*(*C*), which is a normalized quantity that is invariant to a global scaling of *C*’s entries, we can consider

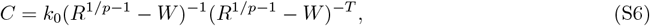

where *W* has been rescaled by some scalar and all scaling has been absorbed into one variable *k*_0_. We shall use this form for all of our following analyses. We will also write *L* = *R*^1−1*/p*^, which differs from the actual neuronal gain in Eq. (S1) by a constant.

First, note that Eq. (3) only holds when the spectral radius *ρ*(*LW*) *<* 1. Using results on the product of isotropic matrices [101, 102], we have

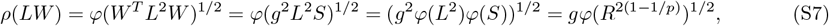

where *S* is a free Poisson element [28] with parameter 1 (i.e., it obeys the Marchenko-Pastur law). Then the condition becomes

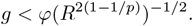

When *p* = 2, we in particular have 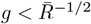 where 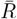 is the population mean firing rate. If we define the normalized parameter 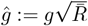, then the condition is simply *ĝ <* 1.

We shall next use tools from operator-valued free probability theory to derive the eigenvalue distribution of the covariance matrix *C* [28] (in particular following pages 245-247 of [28]). Denote by *a, b*, and *c* the corresponding variables in the non-commutative probability space (𝒞, *φ*) of *L*^−1^, *W*, and 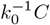 respectively. Note that the notations here are different from those in section B.2 in terms of what *a, b* and *c* represent, but these variables are only used in free probability calculations in each section, so there should not be any danger of confusion and we overload them for readability. *a* is self-adjoint because it is a diagonal matrix with real entries. The circular law states that *b* is a circular element. Eq. (S6) then becomes

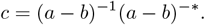

The distribution of *c* is the same as the -2 power of

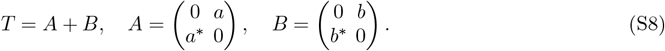

(*b* and −*b* are identical, so we replace *b* with −*b* for simplicity.) *A* and *B* are free 𝒟_2_-valued self-adjoint elements in 𝒜 := *M*_2_(𝒞). Here 𝒟_2_ is the set of diagonal 2 × 2 complex matrices. Explicitly, the conditional expectation *E* : 𝒜 → 𝒟_2_ is defined to be

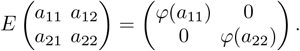

We can take *E* to have a range in 𝒟_2_ because *T* is even due to it having zeros on the diagonal. In particular, *B* is a 𝒟_2_-valued semi-circular element. The covariance function of *B, η* : 𝒟_2_ → 𝒟_2_ is given by

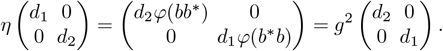

Let *G*_*X*_ be the 𝒟_2_-valued Cauchy transform of *X*. We are ultimately interested in the distribution of *T* ^−2^. Note that there is a relation

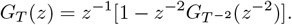

The scalar-valued Cauchy transform *G*_*c*_(*z*), which ultimately determines the eigenvalue distribution *p*_*c*_ of *c*, can be calculated as

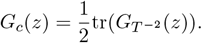

Let us write

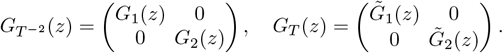

Using results from [28], we have

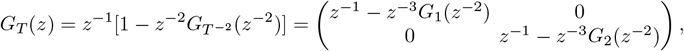

and

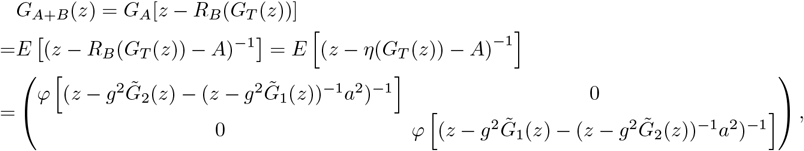

where *R*_*B*_ is the *R*-transform of *B*. Then *T* = *A* + *B* implies that the above two equations are equal, which gives us a system of 2 equations for *G*_1_ and *G*_2_. However, notice that by definition

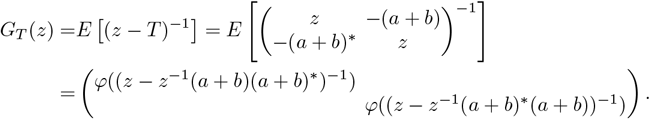

Since (*a* + *b*)(*a* + *b*)^∗^ and (*a* + *b*)^∗^(*a* + *b*) have the same distribution, we know *G*_1_(*z*) = *G*_2_(*z*) = *G*_*c*_(*z*). We therefore get a single equation

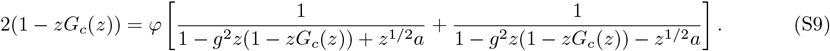

Note that the functional *φ* is with respect to *a*, the variable corresponding to *L*^−1^, which is in turn related to the firing rates *R*. Then we can numerically solve this equation for *G*_*c*_ when we have samples from the firing rate distribution, which in turn gives us the pdf *p*_*c*_. Eq. (S9) was verified by simulation data (Fig. S1b). It is consistent with the result from previous work on uniform gain if we set *a* = 1 [29].

Notice that in our derivation, we only need *L*^−1^ and *W* to be freely independent. If *L*^−1^ is replaced by a positive-definite matrix with a known limiting eigenvalue distribution that is independent of *W*, our derivations can be repeated to arrive at the same result. Conceptually, once there is free independence, we can operate at the level of eigenvalue distributions of *L*^−1^ and *W* and abstract away the element-wise description of the distributions of these matrices. A recent work took a similar approach and constructed *W* from prescribed eigenvalue and eigenvector distributions [103]. Interestingly, they discovered nontrivial dependencies of covariance matrix dimensionality on *W* ‘s spectrum that are hidden in local statistics like motif configurations, highlighting the irreducible value of this approach. It would be interesting to consider connectivity matrices with eigenvalue structures other than the circle law under the free probability framework in future work.

Note that the equation for the Cauchy transform of *C*^−1^ (which is related to *G*_*c*_ through a simple formula) has been derived in previous work using a different method [104].

#### C.3 The effective dimension *D*(*C*)

We next derive expressions for the first few moments of *c* and subsequently the participation ratio. Recall the moment generating function *M*_*c*_(*z*) and its relation with the Cauchy transform:

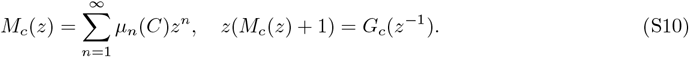

Then Eq. (S9) becomes

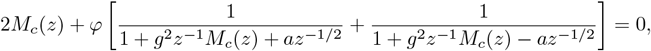

i.e.

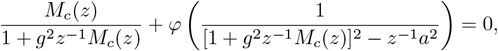

Differentiating with respect to *z* once and twice and setting *z* = 0 gives us the first and second moments of *c*

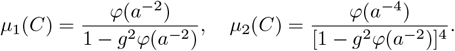

The participation ratio can then be calculated to be

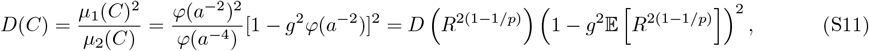

When *p* = 2, this simplifies to Eq. (4). This result also agrees with previous work for uniform gain if we set *R* = *I* [29].

Up to now, we have assumed *C*_0_ = *I* for simplicity. If *C*_0_ is freely independent from *W* and *R*, we can use the same calculation as section B.3 to obtain

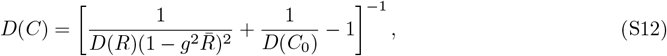

where the positive relation between *D*(*C*) and *D*(*R*) is maintained, and a lower-dimensional *C*_0_ leads to a lower-dimensional *C*. This formula can also be thought of as a generalization of the feedforward formula Eq. (S2) by setting *W* = 0 and *C*_*x*_ = *C*_0_.

#### C.4 The threshold-linear (*p* = 1) transfer function case

When *p* = 1 in Eq. (1), i.e., when the transfer function *f* is threshold-linear [105], *D*(*C*) does not depend on *D*(*R*) if neurons never go completely silent, as the neuronal gains remain constant, with no dependence on firing rates (Fig. S3a-b). However, there may be a dependence if some neurons transition between being inactive and being active across conditions.

Let us consider the simple case of a random subset of *M* neurons being active. Denote the subpopulation of active neurons by the subscript *a*. Then

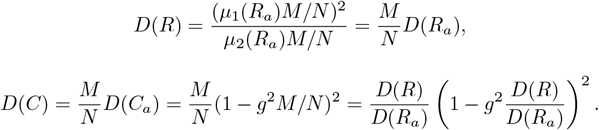

Therefore the relationship between *D*(*C*) and *D*(*R*) depends on how *D*(*R*_*a*_) relates to *M* . Suppose, for instance, that *D*(*R*_*a*_) is constant. Then *D*(*R*) ∝ *M/N* =: *β*. We have

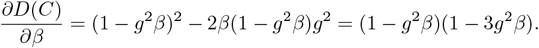

When *g*^2^ ≤ 1*/*3, *D*(*C*) always increases with *β* (and thus *D*(*R*)). When *g*^2^ *>* 1*/*3, *D*(*C*) increases with *β* for *β* ≤ 1*/*(3*g*^2^) but decreases with *β* for *β >* 1*/*(3*g*^2^). We see that the relation between *D*(*C*) and *D*(*R*) depends on the recurrent connectivity diversity *g* and the proportion of active neurons *β*.

#### C.5 A remark on low-rank perturbations and eigenvalue outliers

By “low-rank” perturbations, we mean modifications to the network that result in changes to only a finite number (or more generally, an *O*(*N*^*α*^) number where 0 ≤ *α* < 1) of eigenvalues (in the probabilistic sense). In other words, these effects are measure 0 when considering the eigenvalue distribution in the large *N* limit and therefore cannot be captured by free probability techniques, where the fundamental mathematical object is the eigenspectrum characterized by moments.

There are, however, other techniques one can use to study the effects of low-rank perturbations. For example, Hu & Sompolinsky [29] detailed how to calculate the magnitude of the outlier eigenvalues given the magnitude of the low-rank perturbation along random directions to either *C* (equivalently *C*_0_) or *W* for the case of homogeneous gain. For perturbations to *C*, their technique can be directly transferred to the case of heterogeneous gain we are studying. However, since we do not have an analytic expression for *G*_*c*_ (unlike [29]), the outliers have to be calculated numerically, and it is unclear how much additional insight can be derived compared to directly simulating random matrices. The situation is even more complicated for perturbations to *W*, as the rotational invariance of *C*, which is a crucial component in Hu & Sompolinsky’s derivations, no longer holds in the heterogeneous gain case, and adapting their method to our model may be nontrivial. We therefore leave this to future work.

Perhaps even more importantly, it is usually difficult to simultaneously measure the connectivity *W* or the external input *C*_0_ along with the activity of the same neuronal population. Without access to these information, it is impossible to definitively designate specific eigenvalues of *C* to be due to “bulk” or “low-rank” effects. In our study we identify outlier eigenvalues (see Methods) and obtain the best statistical estimate based on our model as the null distribution. This is sufficient for our purpose here of highlighting the bulk features in *D*(*C*), but a more serious investigation of the low-rank components in the network requires combining types of data other than the neuronal response. For example, there have been recent datasets with large-scale connectivity and activity data in the same population [106]. It would be interesting to study the interaction between low-rank and high-rank components of the network using these dataset, but this is beyond the scope of this paper.

#### C.6 Comparison with previous works

Compared to previous works, our approach is most closely related to that of [29], which also derived the full eigenvalue distribution of the covariance matrix but only for homogeneous gain. They used the replica method instead of techniques from free probability.

Since *D*(*C*) only depends on the first two moments of *C*’s spectrum, which can be equivalently calculated from the first two moments of *C*’s elements (i.e., variances and covariances), methods that focus on the statistics of pairwise (co)variances (instead of the eigenvalue distribution of *C*) can also be employed to study *D*(*C*). Indeed, the formula for *D*(*C*) in a recurrent network with homogeneous gain has also been derived in Dahmen et al. [30] based on the method from [33]. It is important to emphasize that the formula in Dahmen et al. is purported to be more general. Formally, it also reads 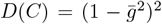, but here 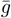 is not just the variance parameter of an i.i.d. connectivity matrix, but the spectral radius of the *effective* connectivity. Then the 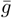 parameter incorporates both the recurrent connection weights and the response properties of neurons, which fits into the broader context of linear response theories of neuronal networks [31, 32]. For example, a recent work conducted a detailed analysis of (co)variance statistics in integrate-and-fire spiking networks [34].

In light of the generality of Dahmen et al.’s result [30], it would be interesting to see if our result can be derived as a consequence of their formula. They used a slightly different expression for the covariance matrix than Eq. (3) that is directly derived from a linear network. To explicitly model how firing rates enter into the effective connectivity through gain and facilitate comparison with our result, we rederive their expression from a nonlinear network with output noise (in contrast to Eq. (S3) which has input noise), namely

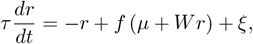

where the trial-to-trial output noise *ξ* has covariance *C*_0_. Again assuming *r* evolves to a steady state in each trial, we have

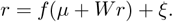

Differentiating the above equation gives us

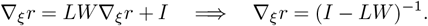

Then

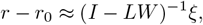

and

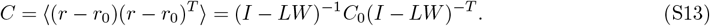

We again consider threshold-quadratic transfer *f* (resulting in *R* = *L*^2^), *C*_0_ = *I*, and i.i.d. Gaussian *W* with mean 0 and variance *g*^2^*/N*. Recall from Eq. (S7) that the spectral radius of the effective connectivity matrix *LW* is 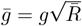, which is also the variance of the elements in *LW*. Then the prediction by Dahmen et al. is

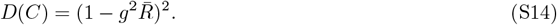

We see immediately that *D*(*C*) only depends on the population average firing rate 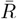 and not on firing rate heterogeneity (i.e., *D*(*R*)). We next derive *D*(*C*) before comparing our prediction with theirs in simulations.

It is less straightforward to study this alternative model using free probability than Eq. (3), because *LW* is no longer a circular element. We instead use an existing formula from [107], where a direct calculation yields the system of equations

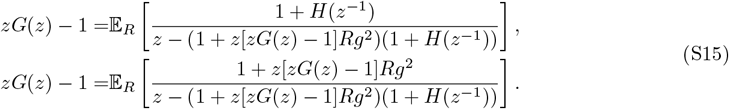

Here, *G*(*z*) is the Cauchy transform of *C, R* is the random variable corresponding to the firing rate distribution (with *R* = *L*^2^), and *H*(*z*) is an auxiliary function that is the integral of *R* against the Stieltjes kernel measure of *C*^−1^. We can solve for *G* and *H* together numerically to obtain the pdf of *C*’s spectrum, the validity of which is verified in simulations (Fig. S2b).

To calculate *D*(*C*), we rewrite the above equations in terms of the moment generating function *M* (*z*) (Eq. (S10)):

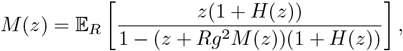

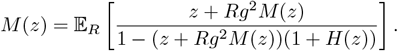

Similar to section C.3, we can differentiate with respect to *z* once and twice and setting *z* = 0 to solve for the first and second moments of *C*:

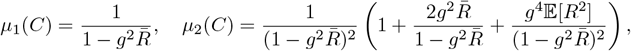

from which we have

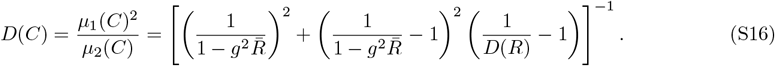

We see that in this formula, *D*(*C*) does depend on *D*(*R*) and agrees with the prediction by Dahmen et al. (Eq. (S14)) only when *D*(*R*) = 1. The validity of Eq. (S16) over Eq. (S14) is verified by simulations (Fig. S2c).

Dahmen et al. also presents a more general formula in their section 4.4.1 that allows *C*_0_ to be a diagonal matrix other than the identity matrix. Notice that taking *C*_0_ = *R* in Eq. (S13) recovers the formula we used for *C* (Eq. (S6) with *p* = 2). However, plugging *C*_0_ = *R* into their formula yields

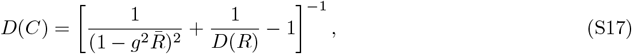

which disagrees with our formula (Eq. (4)). As we have seen, simulations support Eq. (4) over Eq. (S17) (Fig. 3a). Notice that the form of Eq. (S17) can be viewed as combining Eq. (S14) with (S2). This suggests that the formula in Dahmen et al. essentially treats *C*_0_ = *R* as being independent from *LW*, which is not applicable here because *R* = *L*^2^.

While we have demonstrated that the formulae in Dahmen et al. [30] are not directly applicable to capture gain heterogeneity, it is unclear whether their method can be adjusted to correct for the discrepancies we saw here, but this question is beyond the scope of this paper.

### D Temporal and spatial sampling

The theory in the previous sections was derived for the exact covariance matrix of the full network population. In practice, due to constraints of the task and recording time we only have the sample covariance matrix calculated from a finite number of trials or time bins (usually no more than an order of magnitude larger than the number of recorded units), and we only observe a small number of neurons (∼ 10^2^) from a large underlying population (*>* 10^4^). We refer to these limitations as temporal and spatial sampling respectively and shall next examine their implications for the estimate of the effective dimension *D*(*C*). For brevity, we focus on the recurrent network case (as we do in the main text), but similar analyses can be performed for the feedforward network case.

#### D.1 Temporal sampling

The effect of temporal sampling has been studied thoroughly before. Previous work has derived the relation between the covariance eigenspectrum of the original and time-sampled covariance matrix [29, 108], which also applies to Eq. (S9), but we focus on its effect on the effective dimension here. Let *Ĉ* be the sample covariance matrix. Again since *D*(*C*) is normalized, we can without loss of generality assume *k*_0_ = 1. Let *X* ∈ ℝ^*N* × *T*^ be the centered data matrix where each column *X*_*t*_ is the centered spike count vector from one trial, totaling *T* trials. We have

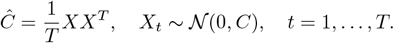

This is equivalent to

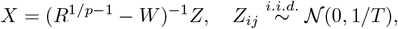

for the recurrent network case. Then

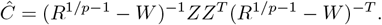

This matrix has the same nonzero eigenvalues as *ČY*, where

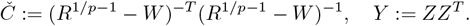

Note that the nonzero eigenvalues of *Č* is again the same as those of *C*, and *Y* is a free Poisson element. Since *Č* and *Y* are independent and *Y* is orthogonally invariant, the product *ČY* is free [28]. Then we can calculate [29]

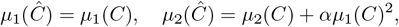

where *α* = *N/T* measures the extent of temporal sampling. Then we have

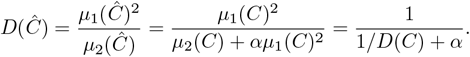

The exact same formula can be derived using the same method for the feedforward network case. A larger *α* (i.e. a smaller number of temporal samples *T*) reduces the effective dimension of the sample covariance matrix *D*(*Ĉ*). Intuitively, a limited amount of samples prevents the system from fully exploring its dimensions.

We can revert the *D*(*Ĉ*) we calculate from data to the original *D*(*C*) by

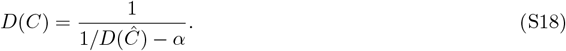

Note that we always normalize the participation ratio by *N* instead of *T*, which is different from the convention used in [29] when *α >* 1.

#### D.2 Spatial sampling

In the uniform gain case, spatial sampling can be formulated as another free product problem [29]. For the feedforward case with heterogeneous gain, this is still possible, as *L* can be absorbed into the spatial sampling matrix (which is essentially a diagonal random matrix with Bernoulli diagonal elements) whose product with *C*_*x*_ is free. One only need to proceed as in [29]. However, it is no longer possible in the recurrent case because *C* is not orthogonally invariant due to heterogeneous gain *L*. However, we can still make a reasonable approximation when the size of the sampled subpopulation is small compared to the whole population. Let *N* = *N*_1_ + *N*_2_ where *N*_1_ is the number of observed neurons and *N*_2_ the unobserved ones. Let *f* := *N*_1_*/N*_2_ and assume 0 *< f* ≪ 1. Recall that after linearization (Eq. (S5)) we have

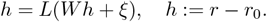

This can be written as

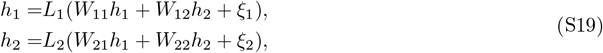

where we have partitioned the vectors and matrices into blocks according to the observed and unobserved population. Using the same notation for the partition, we have that *C*_11_ and *C*_22_ are the covariance matrices of the sampled and unobserved population respectively. If *f* ≪ 1 and the sampling is completely random, we may assume that the term *W*_21_*h*_1_ is much smaller than *W*_22_*h*_2_ and ignore it. That is, we are ignoring the contribution of the sampled population’s input to the unobserved population. Then the covariance matrix of the sampled population is

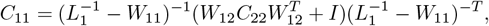

which has the same nonzero eigenvalues as

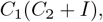

where we have written

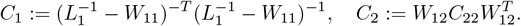

Notice that *C*_1_ has the same nonzero eigenvalues as 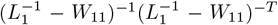, which is just the covariance matrix of the observed population if it were isolated from the rest of the population, and *C*_22_ is just the covariance matrix of the unobserved population when we ignore the input of the observed population (i.e., the *W*_21_*h*_1_ term in (S19)). The behavior of these matrices is known from section C. *C*_2_ has the same nonzero eigenvalues as *C*_22_*D*, where 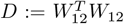. Since *D* is orthogonally invariant and independent of *C*_22_, their product is free. Similarly, the product of *C*_1_ and *C*_2_ + *I* is also free. This allows us to derive a theory for the moments of *C*_11_, although it is an approximation that is only valid when *f* is small.

Recall the formulae for any free variables *a, b*

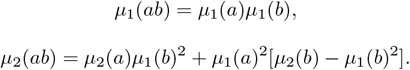

This gives us

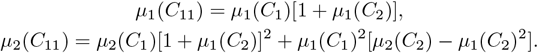

Then

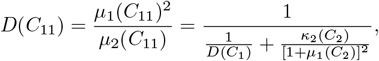

where *κ* denotes the cumulants. For simplicity, we assume *p* = 2 in Eq. (1) in the following. Then

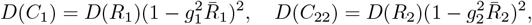

where *g*_1_, *g*_2_ are rescaled versions of *g*. Recall that the elements of *W* are drawn from 𝒩 (0, *g*^2^*/N*). If we want to apply Eq. (4), we need to make sure that the “*g*” here is with respect to the correct population size. That is,

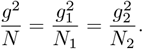

Using *N* = *N*_1_ + *N*_2_, *N*_1_ = *f N*_2_, we have

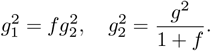

We can also calculate

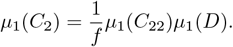

Here the 1*/f* term is to make sure that we only take into account the nonzero eigenvalues of *C*_22_*D. D* is a free Poisson variable with parameter *λ* = 1*/f* with a scaling factor *fg*^2^*/*(1 + *f*). The unscaled free Poisson variable follows the Marchenko-Pastur law and has *µ*_1_ = 1, *κ*_2_ = *λ*. Then

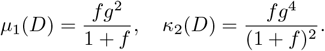

We thus have

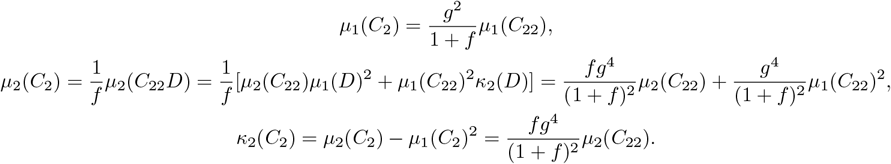

Recall

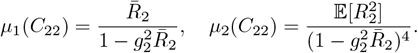

Putting everything together, we have

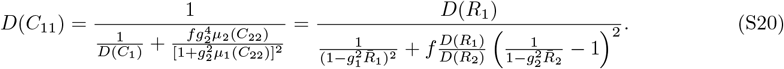

Observe in the denominator that the participation ratios of *R*_1_ and *R*_2_ roughly cancel each other if the sampling is completely random. Then the effective dimension of the observed population is again proportional to its firing rate uniformity, but the coefficient of this relation now depends on the proportion of neurons sampled from the full population as well as the connectivity and activity of the unobserved population which can depend heavily on which neurons are being recorded, making it difficult to interpret the slope of the [*D*(*R*), *D*(*C*)] plot from neural data. As spatial sampling become more drastic, i.e. as *f* gets smaller, the effective dimension *D*(*C*_11_) becomes larger and eventually approaches *D*(*R*_1_) when *f* → 0. (Note that *g*_1_ → 0 as *f* → 0.)

We verified Eq. (S20) using simulation (Fig. S2d). The theory agrees well with simulation when *f* is small, and captures the trend as *f* gets larger. When we record ∼ 100 neurons from an underlying population of *>* 10^4^ neurons, we have *f <* 0.01, which is well within the range where our approximation here is valid.

### E Implications for decoder performance in discrimination tasks

In this section, we explore how firing rate uniformity *D*(*R*) affects neural coding in the context of fine discrimination tasks. Consider two stimulus responses with means *r*_1_, *r*_2_ ∈ ℝ^*N*^ and covariance matrices 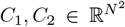 which are related by Eq. (3) (where *R*_i_ = diag(*r*_*i*_), *i* = 1, 2). The mean responses have population average 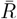 and uniformity *D*(*R*). The discriminability index *d*^′^ is defined by [100]

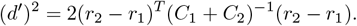

Consider that *r*_1_ and *r*_2_ lie on the simplex

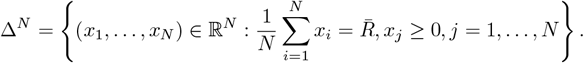

That is, the firing rates are nonnegative and have mean 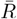. Then the set of points on this simplex with a certain *D*(*R*) is just the intersection of Δ^*N*^ and a (*N* − 1)-sphere of radius 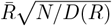 (which we denote with *S*_0_). The intersection *S* := *S*_0_ ∩ Δ^*N*^ is a codimension-2 manifold that is a subset of a (*N* − 2)-sphere of radius 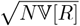, where 𝕍[*R*] is the variance of firing rate across neurons (Fig. 6a).

Let us take a brief detour and look at the topological structure of the manifold *S*. When *D*(*R*) is close to 1, *S*_0_ ⊂ Δ^*N*^ and so *S* = *S*_0_ is just an (*N* − 2)-sphere which is (*N* − 3)-connected. When *D*(*R*) decreases past (*N* −1)*/N*, *S*_0_ intersects with the (*N* −2)-faces of Δ^*N*^, creating a (*N* −2)-dimensional hole and making *S*_0_ (*N* − 4)-connected. As *D*(*R*) gets even smaller, *S*_0_ intersects with faces of Δ^*N*^ that are of lower and lower dimensions. Observe that for the interval of *D*(*R*) where *S*_0_ goes from intersecting the centers of the *M* -faces to intersecting the centers of the (*M* − 1)-faces of Δ^*N*^, the problem of the connectedness of *S* boils down to understanding the situation on each *M* -face, which is again how the intersection of a (*M* − 1)-sphere with the boundaries of a *M* -simplex affects its connectivity. Then it is easy to show by induction that the homotopical connectivity of *S* reduces by 1 each time it starts to intersect with faces of Δ^*N*^ of a lower dimension (beginning at the centers of each face) as *D*(*R*) decreases. The center of an *M* -face of Δ^*N*^ is just the center of mass of *M* + 1 coordinate vertices, which has *M* + 1 equal nonzero coordinates and *N* − *M* − 1 zero coordinates and thus has a *D*(*R*) value of (*M* + 1)*/N* . Then we can summarize

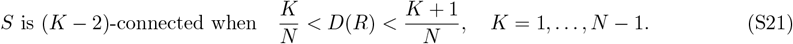

In particular, *S* is disconnected with *N* components when *D*(*R*) *<* 2*/N* . Going back to the problem of discriminating between two stimulus responses, we see that a smaller *D*(*R*) implies both a smaller *D*(*C*) according to Eq. (4) and a larger space *S* to code in, which allows for larger distance *v* := *r*_2_ − *r*_1_ between the stimulus means, both of which are intuitively beneficial to decoding. Another perspective is that given a lower bound on *d*^′^, a smaller *D*(*R*) allows us to pack more stimuli into the space *S* without affecting decoding performance.

Continuing with our assumption that *R* and *W* are independent, we assume that *C* is independent of *v*. Since this is a fine discrimination task, *r*_1_ is close to *r*_2_. In addition, the covariance matrices are smooth functions of the mean responses (Eq. (3)), so *C*_1_ and *C*_2_ must be close as well. We thus assume for simplicity that *C* := *C*_1_ = *C*_2_, but will allow *C*_1_ ≠ *C*_2_ in simulations. Denote the eigenvalue decomposition of *C* as *C* = *U*^*T*^ Λ*U*, where *U* is an orthogonal matrix. Since *C* and *v* are independent, *u* := *Uv* is a completely random rotation of *v* which follows uniform distribution on the sphere of radius |*v*|. Then

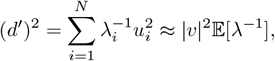

where we used |*u*| = |*v*| . We see that the expression for *d*^′^ factors into two components, the |*v*| ^2^ term depending on the mean *R* and the 𝔼[*λ*^−1^] term depending on the covariance *C*. Let us examine the two factors |*v*|^2^ and 𝔼[*λ*^−1^] separately.

For |*v*| ^2^, since the size of *S* grows as *D*(*R*) becomes smaller, it is reasonable to assume that |*v*| grows linearly with the radius of *S*_0_. Then we have |*v*| ^2^ ∝ 𝕍[*R*] = 1*/D*(*R*) − 1, which increases as *D*(*R*) gets smaller.

For 𝔼[*λ*^−1^], it is the first moment of *C*^−1^. Again for simplicity we only consider the recurrent case here, but the feedforward case can be calculated analogously. Using the same notation as before, we have

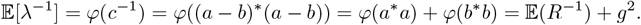

To be more concrete, we assume *r*_1_ and *r*_2_ are drawn from a log-normal distribution ℒ𝒩(*µ, σ*^2^). Recall that for *X* ∼ ℒ𝒩(*µ, σ*^2^), we have

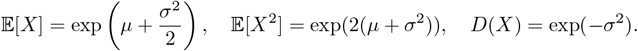

Then

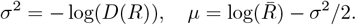

When *R* ∼ ℒ𝒩(*µ, σ*^2^), we have 𝔼(*R*^−1^) = exp (−*µ* + *σ*^2^*/*2) . Then

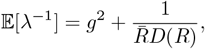

which also increases as *D*(*R*) gets smaller.

Combining the previous sections, we have

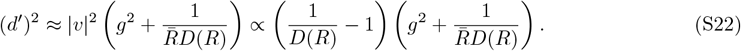

The two factors embody the two ways a decrease in *D*(*R*) can benefit decoding: by enlarging the coding space and lowering the effective dimension of variability. This result is verified in simulations (Fig. 6c).

### F Supplementary figures

**Fig. S1:**
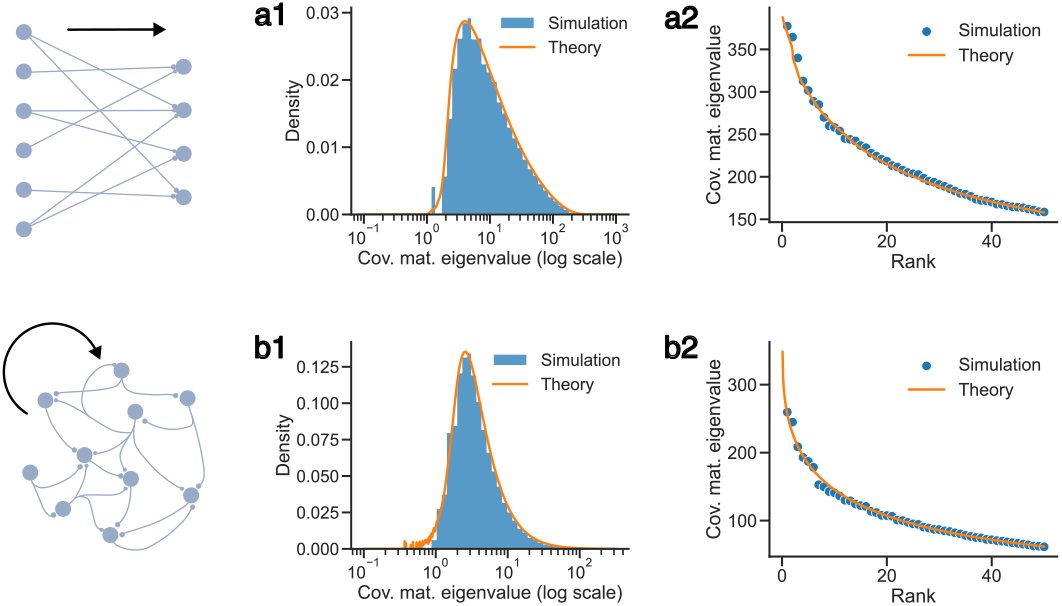
(**a**) Theory of covariance eigenspectrum in a feedforward model in Supplementary Information B.2 verified by simulations. (a1) shows the pdf of eigenvalues and (a2) shows the largest 50 eigenvalues. (**b**) Same as (a), but for the recurrent model in Supplementary Information C.2.

**Fig. S2:**
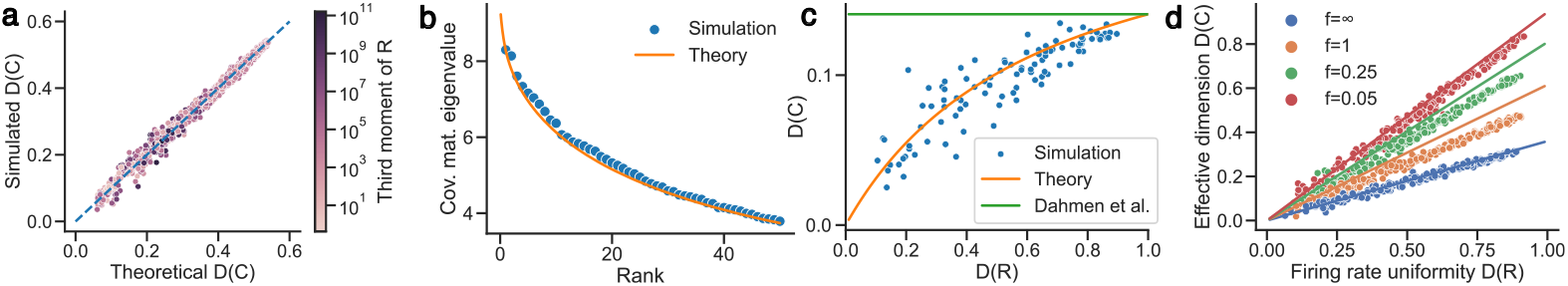
(**a**) Verification that Eq. (4) does not depend on the exact distribution of firing rates *R* beyond *D*(*R*). Here we generated the firing rates *R* by randomly choosing parameters for the generalized gamma distribution [109], which has many common nonnegative continuous distributions as special cases or limiting cases including the gamma distribution, Weibull distribution, the half-normal distribution, and the log-normal distribution which was used in Figs. 1d and 2a. *D*(*C*) is simulated according to Eq. (3) with *g* = 0.15 and 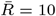. The dashed line is the identity line. It can be seen that our theory predicts *D*(*C*) regardless of the specific distribution of *R* beyond *D*(*R*), which varies dramatically in our simulation as demonstrated by the wide range of third moments of *R*. (**b**) The eigenvalue distribution of *C* as in Eq. (S13) calculated using Eq. (S15) matches simulations. (**c**) Simulation agrees with our formula (Eq. (S16)) instead of that from Dahmen et al. [30] (Eq. (S14)). (**d**) The effect of spatial sampling. The plot shows *D*(*C*) and *D*(*R*) of the recorded population. *f* is the ratio of the size of the recorded population and the unobserved population. *f* = ∞ corresponds to the case where there is no spatial sampling and all neurons in the network are observed. The dots are from simulation and the lines are the theoretical predictions from Eq. (S20). The population size is 1000.

**Fig. S3:**
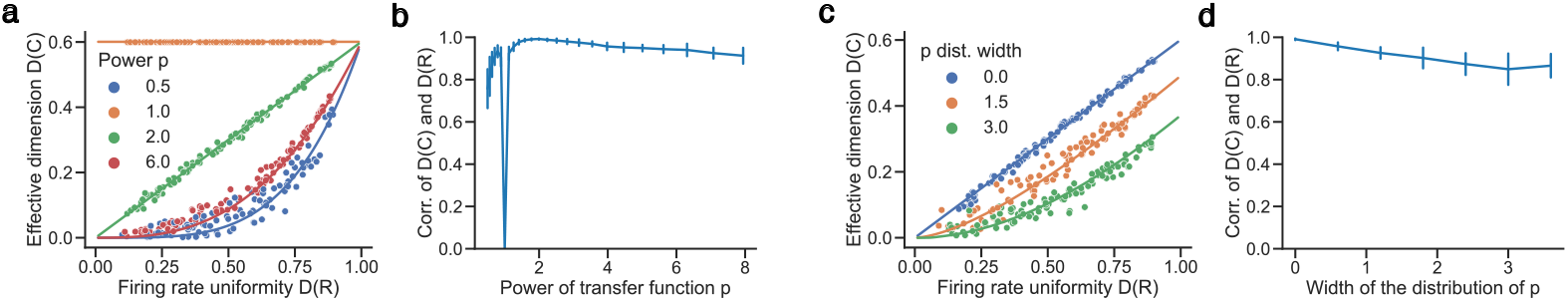
The effect of different transfer functions. All simulations have population size 1000. (**a**) [*D*(*R*), *D*(*C*)] plot when the power *p* in Eq. (1) varies. (**b**) The correlation between *D*(*C*) and *D*(*R*) as *p* varies. *D*(*R*) was sampled in the range [0.3, 0.7] to match neural data. (**c**) Same as (a) but for the heterogeneity of *p*. The power *p* for each neuron is randomly assigned according to a uniform distribution centered at 2 with different widths (meaning the distance between the left and right edges of the uniform distribution). (**d**) Same as (b) but for heterogeneity of *p*.

**Fig. S4:**
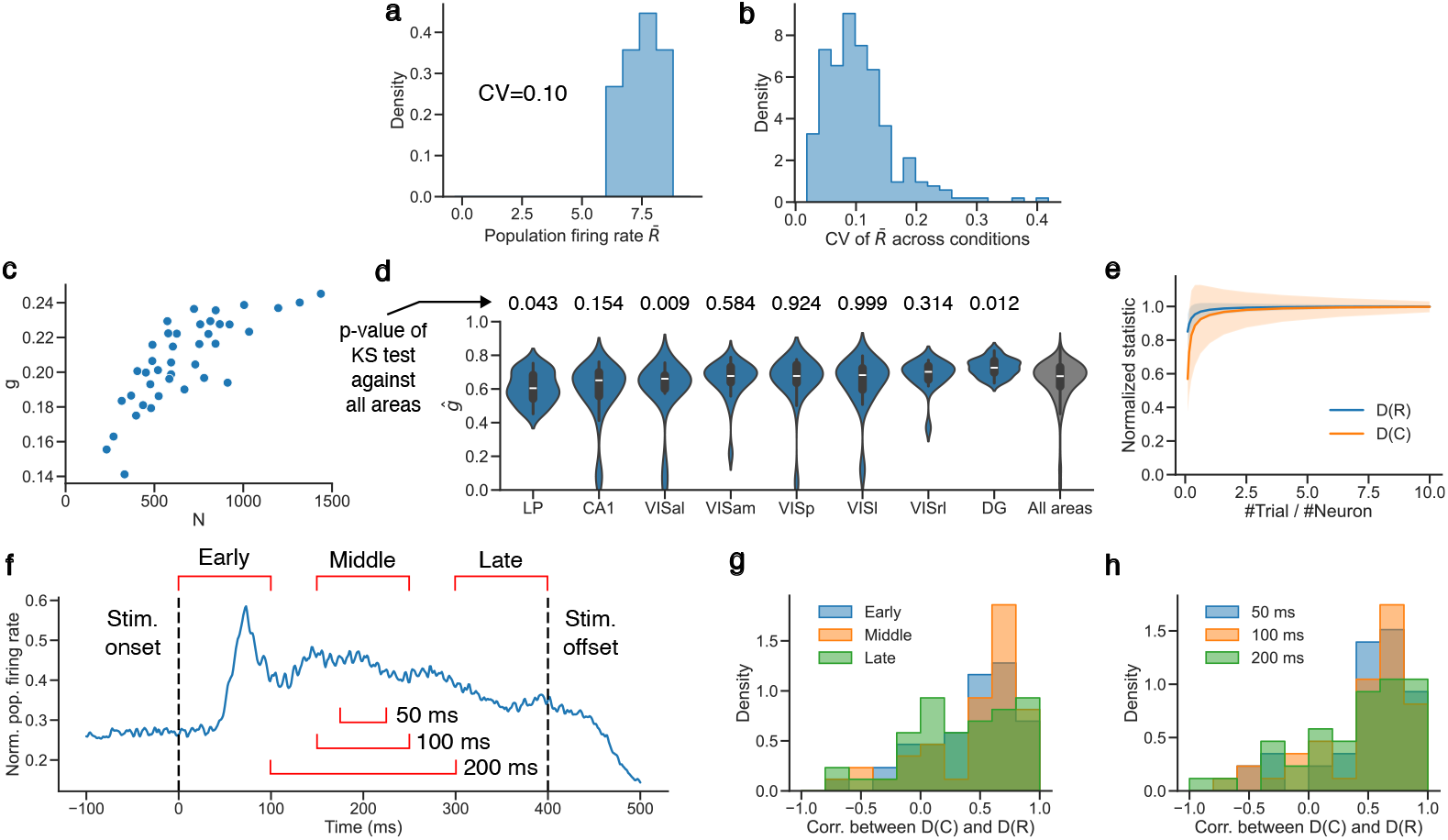
(**a**) The distribution of population average firing rate 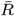 across 16 conditions in a session of monkey LIP recording. It does not vary much, as quantified by a small coefficient of variations (CV). (**b**) The distribution of within-session CV of 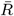 in the same dataset as Fig. 4g [42, 43] across all brain areas and sessions. (**c**) The (*g, N*) pairs discovered by the Bayesian optimization process of monkey LIP data whose cost (Eq. (5)) is within 5% of the optimum. We see that there is a wide distribution of near-optimal solutions to the fitting problem, with a trade-off between the *g* and *N* . (**d**) The distribution of fitted normalized connectivity diversity parameter *ĝ* for each area across sessions in the dataset [42, 43]. Only areas with more than 15 sessions of data are shown. The rightmost plot is the aggregate distribution over all areas and is identical to Fig. 4n. Shown at the top is the p-values of the Kolmogorov–Smirnov (KS) test between each distribution and the overall distribution. Most of the *ĝ* distributions are wide and indistinguishable from the aggregate. (**e**) Empirical estimation of *D*(*R*) and *D*(*C*) as a function of the number of trials compared to population size using the dataset [42, 43]. Note that the *D*(*C*) estimate is already corrected for temporal sampling according to Eq. (6). For each session in each area where the total number of trials is at least 10 times that of the population size, the data is bootstrapped (i.e., trial-subsampled multiple times) to produce the plot. All values are normalized by dividing by the corresponding value when all trials are used. The shaded bands are standard deviations to show the variability in the estimation. (**f-h**) The [*D*(*R*), *D*(*C*)] correlation does not depend significantly on which time window is used in the trial to get spike count data or the window size, demonstrated with the monkey V4 dataset in [40]. (**f**) Normalized trial-averaged population firing rate plot to demonstrate which windows are used. (**g**) Distribution of [*D*(*R*), *D*(*C*)] with different window locations. The median test p-values are: 0.30 (early vs middle), 0.14 (early vs late), and 0.14 (middle vs late). (**h**) Distribution of [*D*(*R*), *D*(*C*)] with different window sizes. The median test p-values are: 0.83 (50 vs 100 ms), 0.53 (50 vs 200 ms), and 0.53 (100 vs 200 ms).

**Fig. S5:**
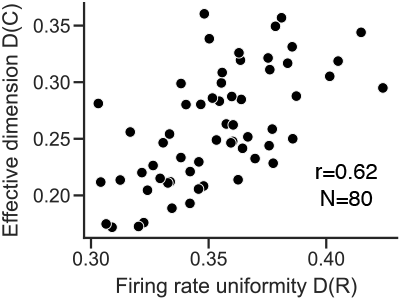
Data recorded from M1 of another human participant as Fig. 4k.

**Fig. S6:**
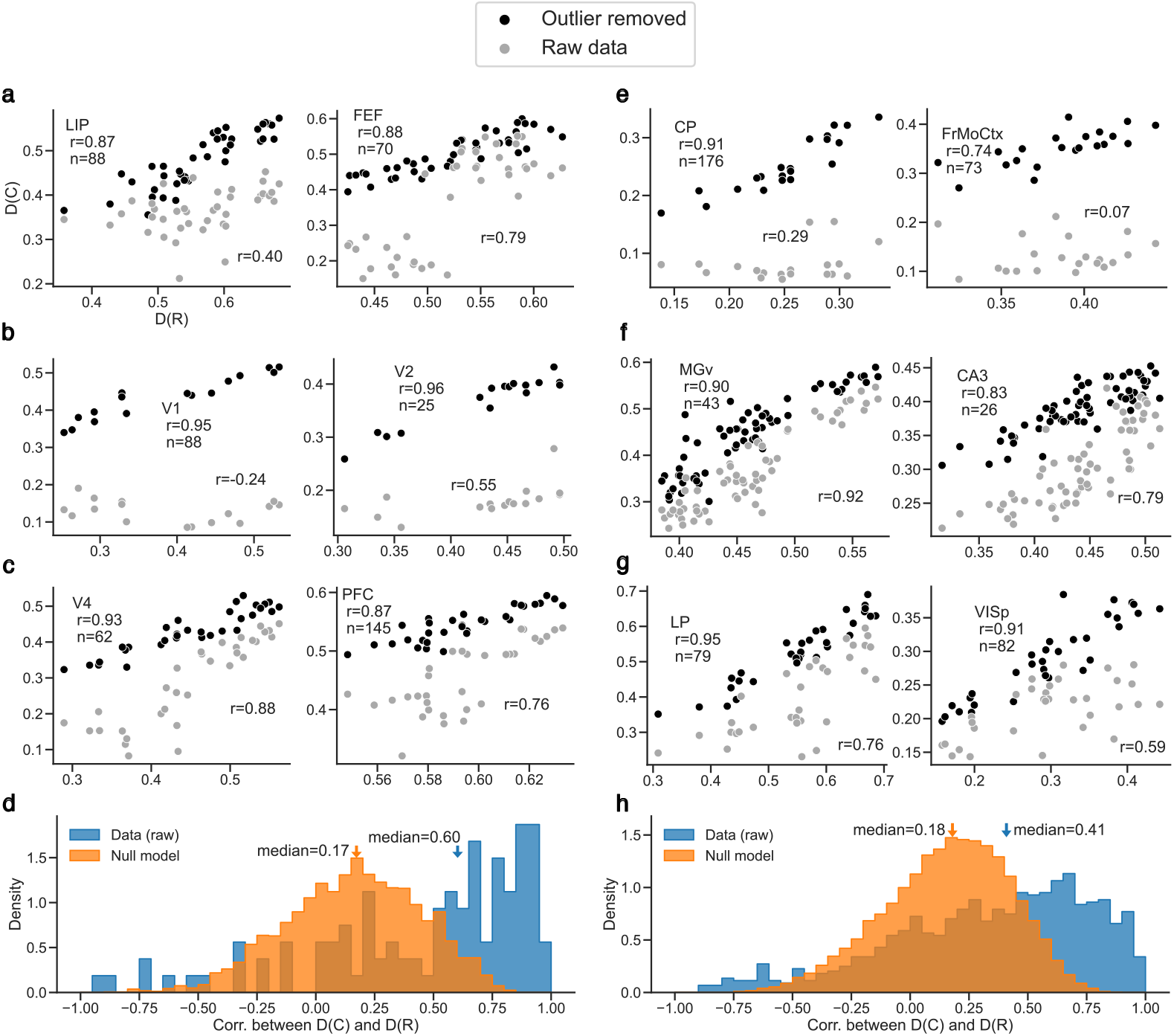
The same as Fig. 4, but compared with the raw data where the outlier eigenvalues are not removed. (a-d) correspond to Fig. 4a-d and (e-h) correspond to Fig. 4f-i. Note that Fig. 4e and j are already the original raw data, and no outlier was identified when producing Fig. 4k. Median test in (d) produces *p* = 1.1 × 10^−7^. Median test in (h) produces *p* = 2.5 × 10^−25^.

**Fig. S7:**
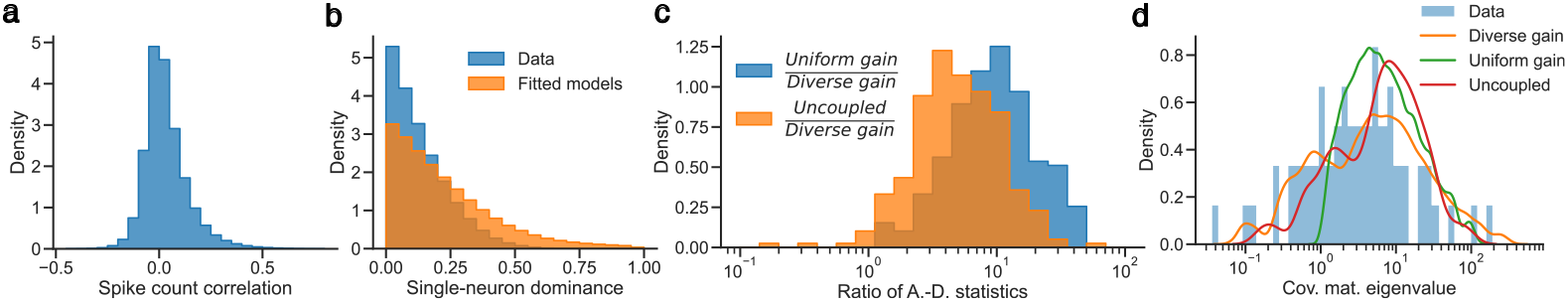
All data in this figure come from dataset [42] and the distributions are over all brain areas and all sessions. (**a**) Distribution of spike count correlation. (**b**) Distribution of the measure for single-neuron dominance of eigenvectors of covariance matrices. Let *v* be an eigenvector and let *v*_1_ and *v*_2_ be the elements of *v* with the largest and second largest absolute values. The measure is defined as (|*v*_1_| − |*v*_2_|)*/* |*v*_1_|, with a value close to one signifying the dominance of one neuron over the rest in this mode and the opposite showing that this mode is shared among multiple neurons. The fact that the histogram concentrates near 0 suggests that there is no correspondence between eigenvectors and neurons, showing that network activity is not a perturbation to uncoupled neurons in neural data. The single-neuron dominance of the fitted models are also shown. The fact that the fitted models (with both heterogeneous weights and gains) have slightly larger single-neuron dominance values suggests that there are additional co-fluctuation patterns present in the data that are not captured by the model, showing that the recorded populations are even further from the uncoupled regime than the model. (**c**) The distribution of the ratios of the Anderson-Darling (A.-D.) statistics between different fitting methods of the covariance matrix eigenvalue distribution. The A.-D. statistics quantifies the quality of the fit, with a smaller value being better (see Methods). It can be seen that our model (i.e. the “diverse gain” model) almost always fits much better than the “uniform gain” model [29] from previous work and the naive “uncoupled” Poisson-like neurons model which simply scales the firing rate distribution to match the eigenvalue distribution, as the ratios are much larger than 1. (**d**) An example showing the performance of different fitting methods.

**Fig. S8:**
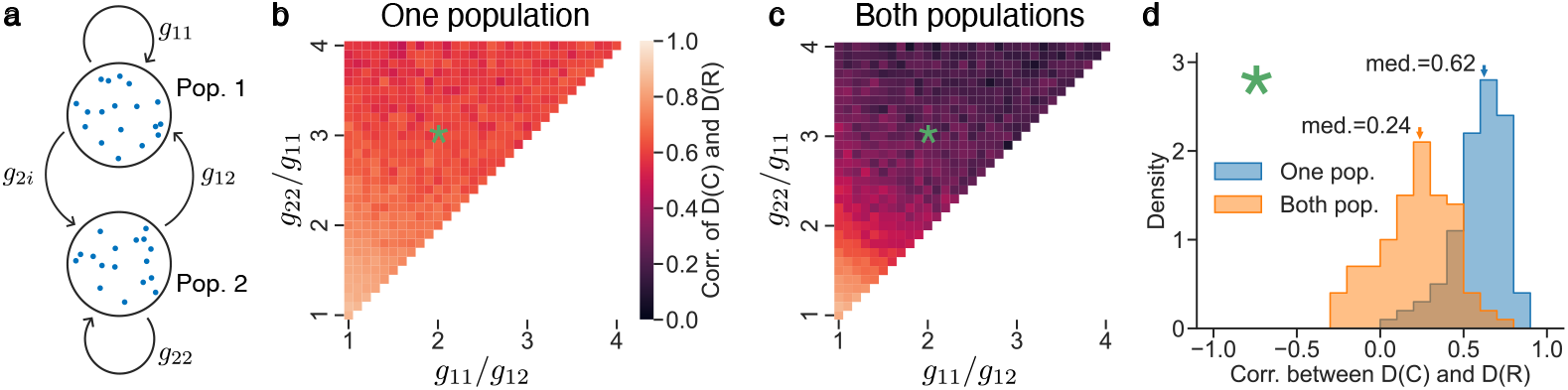
Block structure in the connectivity matrix *W* affects [*D*(*R*), *D*(*C*)] correlation. (**a**) Schematic of the two-population network. Each population has 500 neurons. Connection weights from population *β* to *α* is drawn from a Gaussian distribution with mean 0 and variance 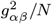. The overall variance of the elements of *W* is held constant (*ĝ* = 0.6) while *g*_22_*/g*_11_ and *g*_11_*/g*_12_ are varied. We also keep *g*_21_ = *g*_12_. To calculate the [*D*(*R*), *D*(*C*)] correlation, 20 *D*(*R*) values was uniformly sampled in the range [0.3, 0.7] to match neural data. This process was repeated 100 times, the averages from which are plotted on the heatmaps. (**b**) The result for each population (averaged across the two populations). (**c**) The result when the activity from the two populations are combined. (**d**) An example distribution of the [*D*(*R*), *D*(*C*)] correlation across realizations of *W* and *R* when *g*_22_*/g*_11_ and *g*_11_*/g*_12_ are at the green asterisk.

**Fig. S9:**
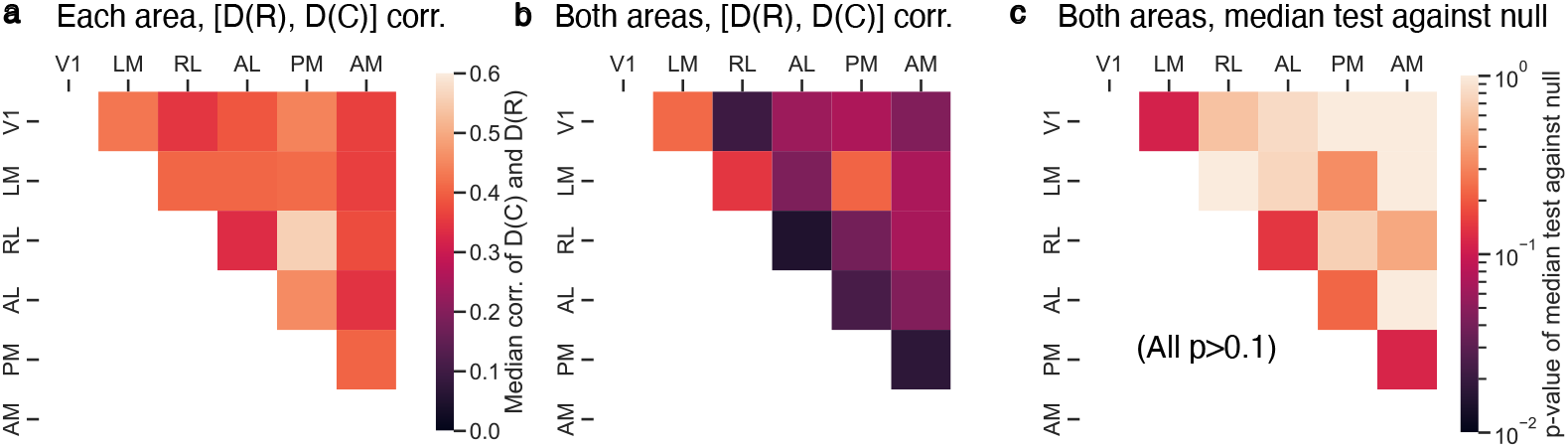
Similar analysis to Fig. 5, but with different two-area combinations in the same dataset as Fig. 5b [42, 43]. The areas are cortical visual areas in mice ordered according to anatomical hierarchy as in [43]. (**a**) Median of each-area [*D*(*R*), *D*(*C*)] correlations across sessions, similar to the yellow violin plots in Fig. 5. (**b**) Median of both-area [*D*(*R*), *D*(*C*)] correlations across sessions, similar to the green violin plots in Fig. 5. (**c**) P-values of the median test between the both-area [*D*(*R*), *D*(*C*)] correlation distribution and the null, similar to the test between the green and blue violin plots in Fig. 5. All both-area [*D*(*R*), *D*(*C*)] correlation distributions are indistinguishable from the null with *p <* 0.1.

**Fig. S10:**
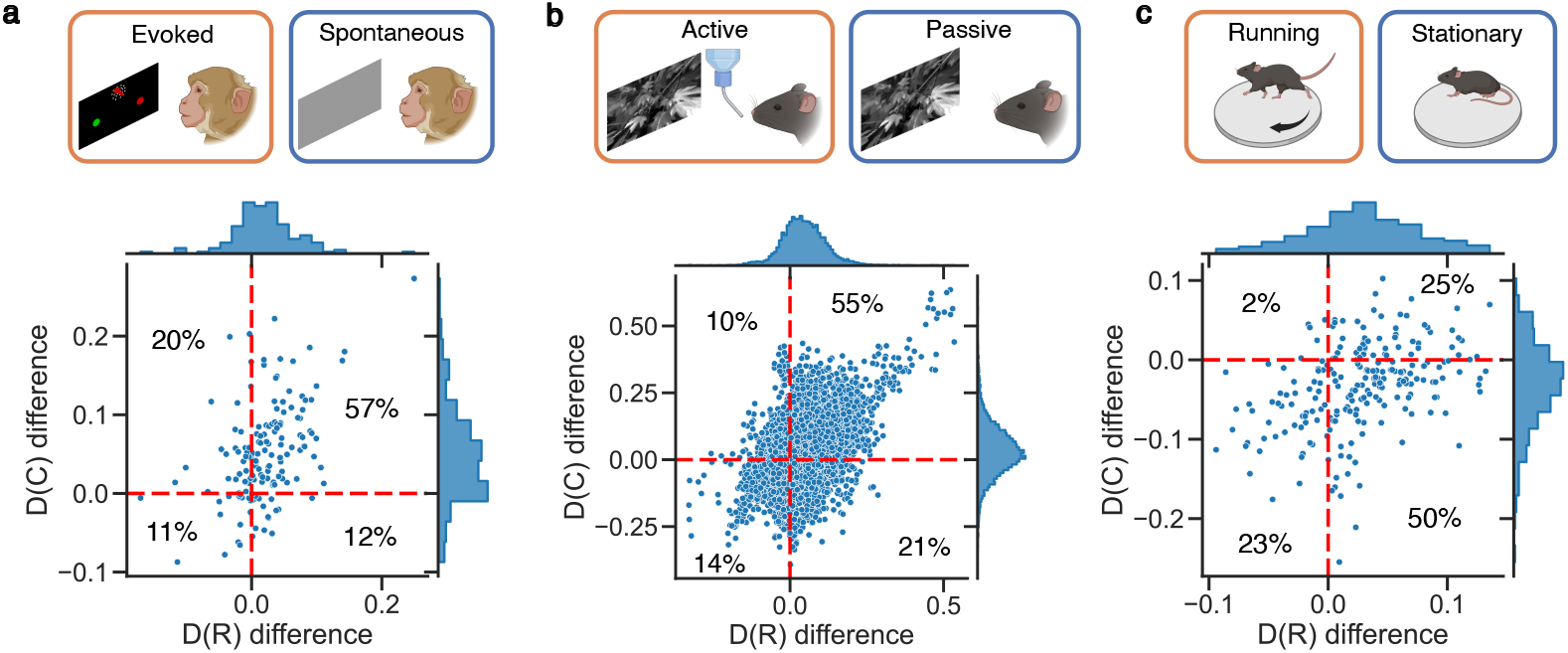
Same as Fig. 7, but without removing the outliers. We see that the most dominant quadrant stays the same as Fig. 7 for (a) and (b), but not (c).

